# Systematic bias in studies of consumer functional responses

**DOI:** 10.1101/2020.08.25.263814

**Authors:** Mark Novak, Daniel B. Stouffer

**Author notes:** Both authors contributed equally to this work. **Corresponding author** Mark Novak, **Email**.

## Abstract

Functional responses are a cornerstone to our understanding of consumer-resource interactions, so how to best describe them using models has been actively debated. Here we focus on the consumer dependence of functional responses to evidence systematic bias in the statistical comparison of functional-response models and the estimation of their parameters. Both forms of bias are universal to nonlinear models (irrespective of consumer dependence) and are rooted in a lack of sufficient replication. Using a large compilation of published datasets, we show that – due to the prevalence of low sample size studies – neither the overall frequency by which alternative models achieve top rank nor the frequency distribution of parameter point estimates should be treated as providing insight into the general form or central tendency of consumer interference. We call for renewed clarity in the varied purposes that motivate the study of functional responses, purposes that can compete with each other in dictating the design, analysis, and interpretation of functional-response experiments.

## Introduction

Functional responses describe how a consumer’s feeding rate responds to its prey’s abundance as determined by behaviour and the composition and conditions of their community and abiotic environment (Abrams & Ginzburg, 2000; Solomon, 1949; Uiterwaal & DeLong, 2020). Applications of the functional-response concept are found in pest control, invasive species management, and conservation biology, and are fundamental to theory in population, community, and evolutionary ecology. Much effort has therefore been devoted to experiments designed to measure and statistically compare the many mathematical models that have been developed to characterize consumer functional responses, with theory evidencing often dramatic differences in the biological inferences and predictions that alternative model forms and parameter values provide (Aldebert & Stouffer, 2018; Arditi & Ginzburg, 2012; Coblentz, 2020; Fussmann & Blasius, 2005).

A dominant focus of the literature on functional responses has been the discussion of so-called prey-, ratio-, and consumer-dependent models (Abrams, 1994; 2015; Abrams & Ginzburg, 2000; Arditi & Ginzburg, 2012; Barraquand, 2014; Tyutyunov & Titova, 2020). These respectively describe feeding rates as responding to prey abundance alone, to the ratio of prey and consumer abundances, or to both prey and consumer abundances more generally. Knowing which of these models best characterize consumer feeding rates can be important because, for example, predictions of population and food-web dynamics are qualitatively altered by either the structural replacement of the classically-assumed prey-dependent Type II model of Holling (1959) for the corresponding ratio-dependent model of Arditi & Ginzburg (1989), or by a parametric change to the strength of mutual consumer effects encapsulated in the consumer-dependent models of Arditi & Akçakaya (1990), Beddington (1975) and DeAngelis *et al.* (1975), and Crowley & Martin (1989).

Although a tremendous amount remains to be learned about consumer functional responses (e.g., Baudrot *et al.*, 2016; Koen-Alonso, 2007; Novak *et al.*, 2017; Uiterwaal & DeLong, 2020), the long-standing debate regarding the relative merits of prey-versus ratio-dependent models in particular is now largely considered by many to be pragmatically resolved: the most recent synthetic assessments of published experiments conclude that most consumer feeding rates respond to both prey and consumer densities, and that they do so in a more general manner than described by exact ratio dependence (DeLong & Vasseur, 2011; Osenberg *et al.*, 1999; Skalski & Gilliam, 2001). More specifically, interference among consumers is considered common, variable in strength, and tending to magnitudes that are “intermediate” to exact prey- and ratio-dependence (Arditi & Ginzburg, 2014; DeLong & Vasseur, 2011; Osenberg *et al.*, 1999). Neither prey- nor ratio-dependence is therefore considered an adequate descriptor of consumer feeding rates (but see Abrams, 2015; Arditi & Ginzburg, 2012; 2014; Tyutyunov & Titova, 2020).

Here we use the perceived resolution of the debate over the consumer-dependent nature of functional responses to bring to light more general issues inherent to the statistical merger of theory and data obtained from functional-response experiments. Our focus is on the use and interpretation of information-theoretic model comparisons (Box 1) and model-specific parameter estimates (Box 2), rather than mathematical analyses of plausibility or the philosophical underpinnings of alternative deterministic model formulations (c.f. Abrams, 2015; Arditi & Ginzburg, 2012; Barraquand, 2014). To assess the extent and severity of these expected statistical issues, we followed prior syntheses of consumer dependence by Hassell & Varley (1969), Hassell (1971), Arditi & Akçakaya (1990), Skalski & Gilliam (2001), and DeLong & Vasseur (2011) and compiled a total of 77 functional-response datasets, incorporating many studies not considered by or available to earlier compilations. Because prior syntheses either used potentially suboptimal statistical methods or included published parameter estimates from across studies using different statistical methods (see summary in Supplementary Materials S2), we re-analyzed all datasets by fitting to them the suite of commonly considered functional-response models and comparing those fits with a now standard information-theoretic approach.

Our analyses reveal systematic bias regarding the inferred density-dependent nature and strength of consumer interference. Bias is driven by variation in experimental sample sizes: studies with smaller sample sizes are biased towards inferences of stronger interference and of particular functional-response model forms. In highlighting the bias for functional-response models of consumer interference, our analyses bring to light the universal but unappreciated bias that small sample sizes introduce to information-theoretic model comparisons and maximum-likelihood parameter estimates for nonlinear functional-response models in general (Boxes 1 & 2). Since all models are wrong (Box, 1976), our primary objective is not to reignite debate regarding prey-, ratio-, and consumer dependence (though we acknowledge this may happen). Rather, we call for clarity and deeper discussion of the varied purposes that motivate the empirical study of consumer functional responses, purposes that can compete with each other in dictating the analysis and interpretation of functional-response experiments.

### Box 1. Bias in information-theoretic model comparisons

Information-theoretic model comparisons have become a mainstay in ecology, as well as in the context of judging the relative performance of alternative functional-response models. The most frequently used information criterion is the Akaike information criterion, *AIC* (Akaike, 1973; 1974). *AIC* estimates the expected relative Kullback-Leibler divergence of a focal model (whose performance in fitting a set of data is being judged) from the “true” model responsible for generating the data. Asymptotically, it minimizes out-of-sample prediction error. A model’s *AIC* reflects the balance between its ability to describe the data as per its likelihood and its complexity as judged by its number of parameters *k*, thereby justifying comparisons of performance for models that differ in complexity. By parsimony, models with smaller *AIC* are deemed to outperform models with larger *AIC*.

However, *AIC* is only an approximately unbiased estimator of the Kullback-Leibler divergence when *k* is small relative to the sample size *n* of the data, and will be negatively biased towards smaller values when *n* is small (Akaike, 1974; Burnham & Anderson, 2002). Sugiura (1978) and Hurvich & Tsai (1989) introduced the corrected Akaike information criterion, *AIC*_*c*_, to correct for this bias by imposing a greater, sample size dependent penalty on a model’s parametric complexity. Because *AIC*_*c*_ converges on *AIC* as sample size increases, it is generally the preferred criterion to use. However, the standard equation for *AIC*_*c*_ was derived in a context of linear regression models with Gaussian errors and remains a biased estimator for nonlinear models and non-Gaussian models, especially when sample sizes are small (Burnham & Anderson, 2002). Even *AIC*_*c*_ may therefore show model-selection bias in favor of more complex functional-response models when sample sizes are small.

Furthermore, a model’s complexity is not just a function of its number of parameters but also of its functional form. Two models with an equal number of parameters can combine these in different ways, leading to differing degrees of flexibility (Gutenkunst *et al.*, 2007; Ly *et al.*, 2017). This form of complexity (a.k.a. geometric complexity) may be considered problematic in the context of assessing model performance because a model with greater flexibility will have larger estimation uncertainty associated with each of its parameters’ point estimates that together “conspire to cancel each other out for the special purpose of fitting the observed data” (Myung, 2000). Criteria such as *AIC* and *AIC*_*c*_ that use only the maximum-likelihood point estimates ignore this flexibility, the effect of which diminishes with increasing sample size for alternative criteria that do consider it (Myung, 2000; Myung *et al.*, 2006). Multiplicative models models like the Arditi-Akçakaya model are generally expected to exhibit greater flexibility than additive models like the Beddington-DeAngelis model (Fig. 1, see Supplementary Materials S3), hence are expected to be preferentially selected by *AIC* and *AIC*_*c*_ when sample sizes are small.

**Figure 1:**
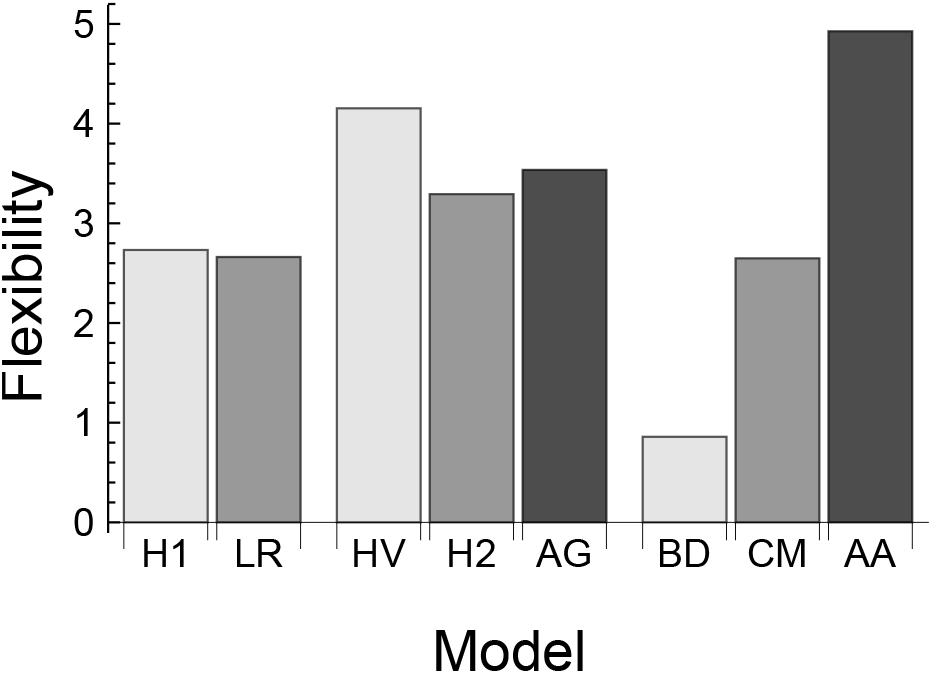
Model complexity as assessed by the flexibility term of the Fisher Information Approximation applied to each of the functional-response models we considered (Table 1). We assumed an experimental design entailing *N* ∈ {2, 4, 8, 16, 32} and *P* ∈ {1, 2, 5} treatment levels with prey replacement, and that the numbers of prey eaten were samples from a Poisson distribution constrained such that no more than an expected 32 prey were eaten in any treatment level in a single unit time period *T*. Note that flexibility will differ for different experimental designs and may only be compared among models having the same number of parameters; an additional term in the Fisher Information Approximation penalizes models for their number of parameters (see Supplementary Materials S3).

### Box 2. Bias in maximum-likelihood parameter estimates

The sample size dependent bias of maximum-likelihood parameters estimates for nonlinear models differs from the model-comparison bias discussed in Box 1. We illustrate the general problem using a simple example co-opted from Box (1971), then show its relevance to the estimation of functional-response parameters.

Box (1971) considered the issue of estimating the rate parameter *θ* for a model of exponential decay,

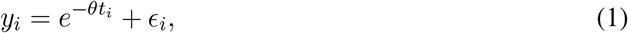

where all *i* = 1, 2, . . ., *n* observations of response variable *y*_*i*_ are made at the same time (*t*\* = *t*_1_ = *t*_2_ = ... = *t*_*n*_) and the є_*i*_ residuals are assumed to be independent and described by a Gaussian distribution with a mean of zero and a variance of *σ*^2^. The value of *θ* that maximizes the likelihood of the data is

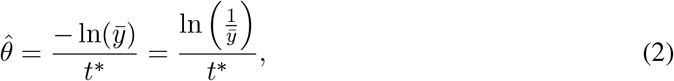

where 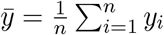 is the arithmetic mean of all observations. Since all observations were taken at the same time, *ȳ* will follow a Gaussian distribution whose mean of exp(−*θt*\*) will have a standard error of 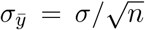. Because the numerator of eqn. 2 is a concave-up function of *ȳ*, Jensen’s inequality dictates that 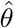 will be biased upward with respect to the true value of *θ* unless *σ*_*ȳ*_ = 0. Estimation bias is therefore an intrinsic consequence of observation noise or variation in the assumed decay process, but becomes smaller as sample size increases.

Equivalent variation-induced bias occurs for the parameter estimates of all nonlinear functional response models (see Supplementary Materials S7 for details and derivations of qualitative bias in representative models). For example, for the Arditi-Akçakaya model on which most prior inference of consumer interference has focused, the maximum-likelihood estimator for the interference strength *m* when eaten prey are replaced is

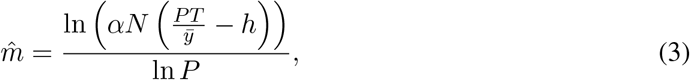

where *ȳ* is the arithmetic mean number of eaten prey and all other parameters are defined in Table 1. Because *ȳ* cannot exceed the possible number of prey eaten, *PT/h*, the numerator of eqn. 3 is positive and is also a concave-up function of *ȳ*. This makes 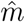 biased upward with respect to the true value of *m* unless there is no residual variation among experimental replicates or sample sizes are large, regardless of whether the numbers of prey eaten in the course of an experiment are assumed to be samples from a Poisson or Gaussian distribution. The estimator for *m* is similarly biased when the proportions of prey eaten are assumed to be drawn from a binomial distribution for non-replacement studies. In fact, for non-replacement studies, even the estimator for the attack rate of linear functional response models is biased. Since maximum-likelihood estimators derived assuming a Gaussian likelihood are equivalent to the estimators of least squares regression, bias will occur when applying the latter as well. Further, since maximum-likelihood estimators derived under a log-normal likelihood are a function of the geometric mean *y* rather than the arithmetic mean, and the geometric mean will always be less than the arithmetic mean, estimation bias for parameters like *m* will be compounded when using regression on log-transformed counts of prey eaten.

**Table 1:** The eight deterministic functional-response models we considered for describing the rate at which prey (hosts) are eaten (parasitized). Variables *N* and *P* refer to the abundances of prey (hosts) and predators (parasitoids). Parameters *a* and *α* respectively refer to the attack rates of the Holling- and ratio-like models; their dimensions differ (Arditi & Ginzburg, 2012). Parameters *c* and *m* respectively represent interference strengths for Holling- and ratio-like models; we use separate symbols because their dimensions also differ. Note that *m* is constrained to positive values in the formulations used here; much previous work uses equivalent formulations in which *m* is negative or is made positive by explicit negation. When fitting the BD and CM models, we replaced *P* with *P* 1 for datasets in which *P* reflected a count rather than density. The number of parameters, *k*, refers to the number of free parameters in each model because only these determine the mean and variance of the Poisson and binomial statistical models we employed. For replacement studies to which the Poisson was applied, the predicted count of prey eaten corresponds to the functional response multiplied by *P* and the time period of the experiment or observation window, *T*. For non-replacement studies to which the binomial was applied, the predicted proportion of prey eaten by *P* consumers after time period *T* was solved for analytically (see Supplementary Materials S7).

| Name | Abbreviation | Model | Parameters ($k$) |
| --- | --- | --- | --- |
| Holling Type I | H1 | $aN$ | 1 |
| Holling Type II | H2 | $\frac{aN}{1+ahN}$ | 2 |
| Beddington-DeAngelis | BD | $\frac{aN}{1+ahN+cP}$ | 3 |
| Crowley-Martin | CM | $\frac{aN}{(1+ahN)(1+cP)}$ | 3 |
| Linear Ratio-dependent | LR | $\frac{\alpha N}{P}$ | 1 |
| Arditi-Ginzburg | AG | $\frac{\alpha N}{P+\alpha hN}$ | 2 |
| Hassell-Varley | HV | $\frac{\alpha N}{P^m}$ | 2 |
| Arditi-Akçakaya | AA | $\frac{\alpha N}{P^m+\alpha hN}$ | 3 |

## Methods

### Data compilation and functional-response models

We attempted to consider all published functional-response datasets of predator and parasitoid consumers having variation in both prey (host) and consumer abundances, the majority of which represented manipulative experiments (Table S1). Potential datasets were identified in three ways:

1. by assessing the data sources of the above-referenced prior syntheses;
2. by assessing all publications citing the above-referenced prior syntheses, as well as Abrams & Ginzburg (2000); and
3. by cross-referencing our compiled list of datasets with the more extensive, primarily prey-only functional-response dataset compilations of Rall *et al.* (2012) and DeLong & Uiterwaal (2018).

We stopped compiling datasets in July of 2018. When possible we obtained the original data from the authors. Otherwise we extracted data points, or means and associated uncertainty estimates, from the publication, either from provided tables or using GraphClick (2010). A few published studies were not included in our analyses because, for example, prey experienced significant mortality not attributable to predation (e.g., Schenk *et al.*, 2005; Stier *et al.*, 2013; Stier & White, 2014) or because observations were not all independent (e.g., Blowes *et al.*, 2017; Cresswell, 1998), which necessitated tailored models beyond the scope of our analyses. Similarly, a few studies in which prey abundances or the numbers of prey eaten were quantified as non-integer values (i.e. densities or feeding rates, e.g., Hebblewhite 2013) were not included since these necessitated alternative statistical models, though for most such previously analyzed studies these data could be converted to counts (i.e. numbers of prey available and eaten in a given amount of time, e.g., Vucetich *et al.* (2002)). Unlike some prior syntheses (Arditi, *pers. comm.*; see also Abrams, 2015), we did not exclude datasets in which significant prey-depletion occurred (but see Fig. S1). Datasets entailing consumers feeding on multiple prey species were included and treated as separate prey-specific datasets only when this was done by the original authors (e.g., Chan *et al.*, 2017).

We considered a total of eight commonly considered deterministic functional-response models (Table 1): the prey-dependent Holling Type I and Type II forms (Holling, 1959), the corresponding ratio-dependent forms (Arditi & Ginzburg, 1989; Sutherland, 1983), and the more general consumer-dependent forms of Beddington (1975) and DeAngelis *et al.* (1975), Crowley & Martin (1989), Hassell & Varley (1969), and Arditi & Akçakaya (1990). Of the two Holling-like consumer-dependent models, the Beddington-DeAngelis model assumes interference occurs only when consumers are searching for prey, while the Crowley-Martin model assumes it occurs equally during both searching and handling. Of the two ratio-like consumer-dependent models, the Hassell-Varley model assumes that handling times are inconsequential and the Arditi-Akçakaya model does not; both assume that interference among consumers occurs through a generic multiplicative process. We did not consider any of the many additional nor less-phenomenological functional-response models that have been proposed (e.g., Abrams, 2010; Jeschke *et al.*, 2002), including parasitoid-specific models (e.g., Rogers, 1972). Hence, for some parasitoid datasets and models, attack rate estimates are better interpreted as reflecting encounter rates (Arditi, 1983).

### Model-fitting and performance

We fit all functional-response models to each dataset by first obtaining maximum-likelihood parameter estimates using the *sbplx* optimization function of the *nloptr* R package (Johnson, 2020), then passing these to the *mle2* function of the *bbmle* package (Bolker, 2020) using the default optimizer to estimate parameter uncertainty. For each dataset, we assumed one of two statistical models: a Poisson likelihood for studies in which eaten prey were continually replaced, and a binomial likelihood for studies in which they were not. For non-replacement studies, we generated predictions for the number of eaten prey using the Lambert W function (Lehtonen, 2016) after confirming that numerical integration using *odeintr:rk4* (Keitt, 2017) gave equivalent results. All parameters were constrained to be positive by exponentiation (Bolker, 2008). For observational field studies, we followed the original author’s assumed prey replacement scenario (Table S1). Parasitoids were classified as either discriminatory or non-discriminatory, with studies of non-discriminatory parasitoids being treated as replacement studies (Table S1). Data available only in the form of means and uncertainties were analyzed by a parametric bootstrapping procedure in which 250 new datasets were created assuming either a treatment-specific binomial or Poisson process as dictated by the study’s replacement of prey. More specifically, for a treatment level of *N* prey having *n* replicates and a mean *μ* and standard error *s* prey eaten, we drew *n* random values *x* from a Gaussian distribution, 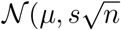, to parameterize a Poisson distribution (setting *x <* 0 to *x* = 0) or binomial distribution (setting *x <* 0 to *x* = 0 and *x > N* to *x* = *N*) to generate integer counts of prey eaten. We considered the median of the resulting distribution of parameter estimates as the point estimate. In one case where no estimates of uncertainty were available (Katz, 1985), we treated the available means as raw data, as done in prior syntheses.

We assessed the relative performance of the fitted models for each dataset by ranking them in two ways: by their *AIC*_*c*_ and by their mean absolute deviation (*MAD*). Despite its assumptions (Box 1), *AIC*_*c*_ is typically the preferred information criterion when sample sizes are small or vary across datasets; it converges on the most widely-used information criterion, *AIC*, as sample size increases. As with *AIC*, models having *AIC*_*c*_ values within 2 units of the best-performing model are considered to have equivalent support (Burnham & Anderson, 2002). In theory, *AIC* is asymptotically optimal in the sense of minimizing a model’s out-of-sample mean squared error (*MSE*). *MAD* is similar to *MSE* but describes the within-sample differences between observations and predictions, and is more easily interpreted by not differentialy re-scaling deviations of different magnitude the way that squaring deviations does. For the additional purpose of judging model performance in a relative goodness-of-fit sense, we also evaluated each model’s *MAD* as a percentage of the *MAD* of the best-performing model having the lowest *MAD*, arbitrarily considering a 1% difference as reflecting a negligible difference in model performance. We then looked for systematic variation across all datasets in their rankings of alternative models by ordering datasets by their sample size. Sample size was defined as the total number of experimental replicates (within and across treatments) or observations (e.g., time points) as defined by the original authors.

### Parameter inference

Recent synthetic assessments of consumer dependence fall into two categories: those that evaluated relative support for alternative model forms (as above, e.g., Skalski & Gilliam, 2001), and those that inves-tigated magnitudes of interference as estimated using the consumer-dependent Arditi-Akçakaya model (e.g., Arditi & Ginzburg, 2012; DeLong & Vasseur, 2011). The Arditi-Akçakaya model combines the saturating form of the Holling Type 2 and Arditi-Ginzburg models with the multiplicative interference effects of the Hassell-Varley model. Its multiplicative nature makes the interference strength parameter, *m*, dimensionless (unlike all other parameters of this and all other models). This permits direct comparisons across studies, with estimates of *m* = 0 evidencing perfect prey dependence and estimates of *m* = 1 evidencing perfect ratio dependence. Therefore, and although bias due to low sample size may occur for the parameters of all nonlinear models (Box 2, Supplementary Materials S7), we focused our assessment of bias on *m*, plotting estimates as a function of each study’s sample size and other covariates (i.e. predator or parasitoid, replacement or non-replacement, no depletion or depletion, raw or bootstrapped data). To assess the specificity of inferences to our chosen statistical approach, we also repeated our estimation of *m* by employing the method used by Arditi & Akçakaya (1990) for all datasets where this method could be applied (see Supplementary Materials S5).

Past assessments of interference strengths have relied on point estimates alone, just as is implicitly done when model performance is assessed using information criteria such as *AIC*_*c*_ (Box 1). However, assessing the uncertainty of parameter estimates is equally important when drawing inferences (Aldebert & Stouffer, 2018; Osenberg *et al.*, 1999). We approximated standard errors in one of three ways: For non-bootstrapped datasets, we first attempted the *confint* profiling method of *bbmle* (Bolker, 2020) assisted by the *SEfromHessian* function of the *HelpersMG* package (Girondot, 2020). On occasion, this failed because the optimization converged but the likelihood surface was nearly flat around the optimum. In these cases, we resorted to the quadratic approximation of *mle2* (Bolker, 2020). For bootstrapped datasets, we used the 16^*th*^ and 84^*th*^ quantiles of the distribution of estimates, corresponding to ±1 standard deviation of a normal distribution (Efron & Tibshirani, 1994).

## Results

### Model performance

As judged by *AIC*_*c*_, the Crowley-Martin, Arditi-Akçakaya and Arditi-Ginzburg models were respectively each ranked first by 23 (29.9%), 17 (22.1%) and 17 (22.1%) of the 77 datasets (Fig. 2A, Table 2). This was followed by the Beddington-DeAngelis model which ranked first for 11 (14.3%) datasets. No datasets ranked the linear ratio-dependent model as first. The Beddington-DeAngelis and Arditi-Akçakaya models were the most frequent models to be ranked second at a respective 35 (45.5%) and 20 (26.0%) times. In total, 46 (59.7%) datasets had a single model outperform all others by more than two *AIC*_*c*_ units. For the 23 datasets where the Crowley-Martin model was ranked first, the Beddington-DeAngelis, Arditi-Akçakaya, and Arditi-Ginzburg models were each within two *AIC*_*c*_ units for one dataset. For the 17 datasets where the Arditi-Akçakaya model was ranked first, at least one of three models – the Beddington-DeAngelis, the Crowley-Martin, and the Arditi-Ginzburg model – were within two *AIC*_*c*_ units a total of 6 (35.3%) times.

**Figure 2:**
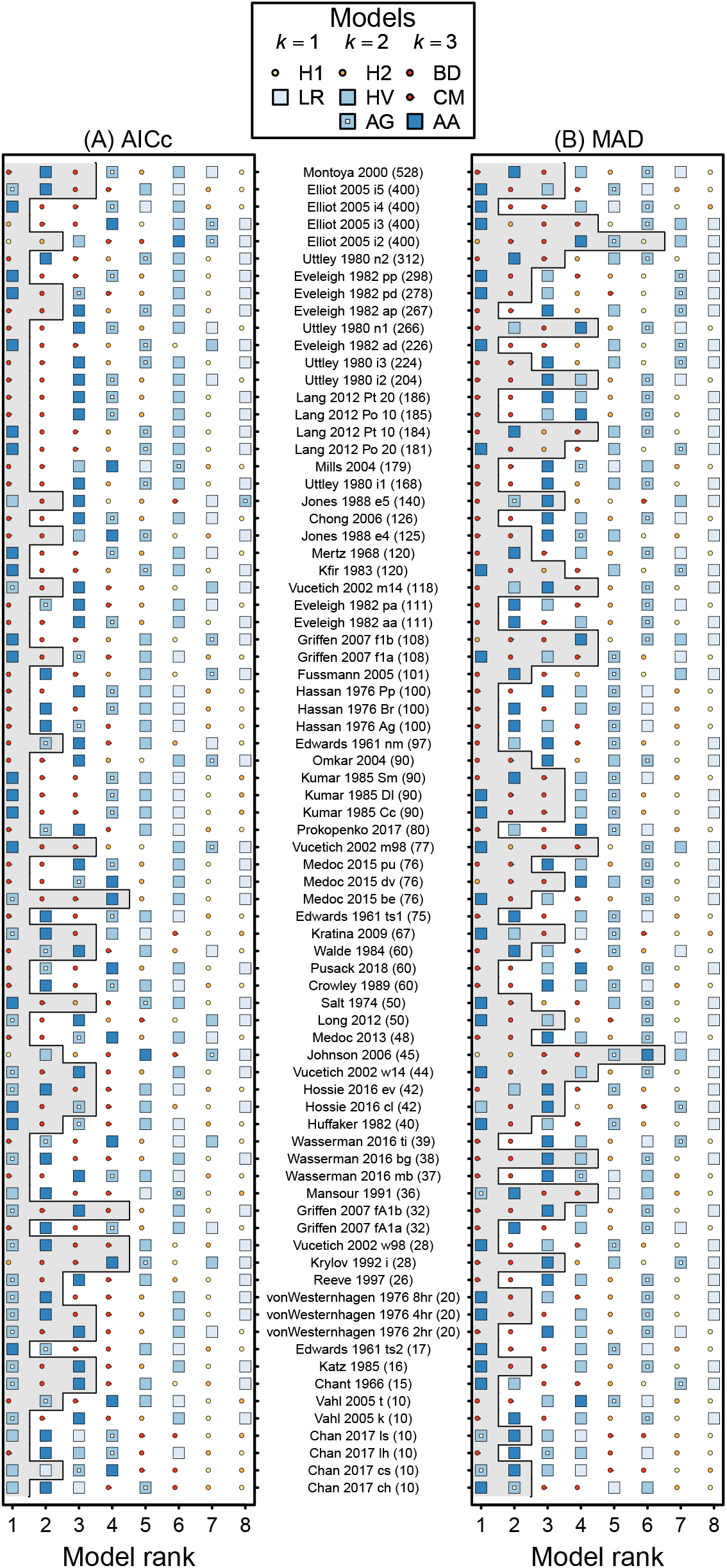
The rank-order performance of functional-response models as judged by (A) *AIC*_*c*_ and (B) *MAD*, with datasets ordered by their sample size. The gray region in (A) demarcates models interpreted as having equivalent support to the top model (i.e. Δ*AIC*_*c*_ < 2). The gray region in (B) demarcates models having a *MAD* value within 1% of that of the top-ranked model. See Table 1 for model abbreviations.

**Table 2:** The number of datasets for which each functional response model achieved a given rank relative to all other models as judged by *AIC*_*c*_ or by *MAD*. Models having equivalent support to the top-ranked model are indicated in Fig. effig:Ranks.

| Rank | H1 | H2 | BD | CM | LR | HV | AG | AA |
| --- | --- | --- | --- | --- | --- | --- | --- | --- |
| *AIC<sub>c</sub>* |  |  |  |  |  |  |  |  |
| 1 | 2 | 2 | 10 | 23 | 0 | 5 | 17 | 18 |
| 2 | 0 | 1 | 35 | 12 | 1 | 1 | 8 | 19 |
| 3 | 0 | 2 | 18 | 16 | 2 | 4 | 8 | 27 |
| 4 | 1 | 13 | 9 | 20 | 0 | 1 | 22 | 11 |
| 5 | 3 | 28 | 2 | 3 | 4 | 24 | 12 | 1 |
| 6 | 9 | 4 | 3 | 3 | 18 | 37 | 2 | 1 |
| 7 | 34 | 19 | 0 | 0 | 12 | 5 | 7 | 0 |
| 8 | 28 | 8 | 0 | 0 | 40 | 0 | 1 | 0 |
| *MAD* |  |  |  |  |  |  |  |  |
| 1 | 1 | 3 | 18 | 26 | 0 | 3 | 1 | 25 |
| 2 | 0 | 3 | 34 | 13 | 0 | 2 | 7 | 18 |
| 3 | 0 | 4 | 18 | 12 | 0 | 1 | 17 | 25 |
| 4 | 1 | 11 | 4 | 19 | 3 | 3 | 28 | 8 |
| 5 | 4 | 28 | 1 | 4 | 5 | 22 | 13 | 0 |
| 6 | 12 | 8 | 2 | 3 | 12 | 35 | 4 | 1 |
| 7 | 31 | 17 | 0 | 0 | 11 | 11 | 7 | 0 |
| 8 | 28 | 3 | 0 | 0 | 46 | 0 | 0 | 0 |

Sample sizes varied greatly among the datasets and were skewed towards smaller studies (range = 10–528, median = 80, mean = 115.1, Fig. 2). The smaller their sample size, the greater was the proportion of datasets that ranked the Arditi-Ginzburg, Hassell-Varley, or Holling Type II models as first (Fig. 3A). Thus, for example, for the 39 datasets having sample sizes equal to the median sample size or less, the Arditi-Ginzburg model was ranked first 15 (38.5%) times while the Crowley-Martin, Arditi-Akçakaya, and Beddington-DeAngelis models were ranked first a respective 12 (30.8%), 5 (12.8%), and 1 (2.6%) times, and were ranked second a respective 3 (7.7%), 13 (33.3%) and 15 (38.5%) times. Correspondingly, the more that datasets with larger sample sizes were considered, the greater was the proportion that ranked either the Crowley-Martin, Arditi-Akçakaya, or Beddington-DeAngelis models as first (Fig. 3A), and the Beddington-DeAngelis model as second (Fig. 3B). Datasets with larger sample sizes were somewhat better able to discriminate among models than were smaller datasets, with 19 (48.7%) of the 39 smaller datasets and 27 (71.1%) of the 38 larger datasets having a single model outperform all others by more than two *AIC*_*c*_ units (Fig. 2).

**Figure 3:**
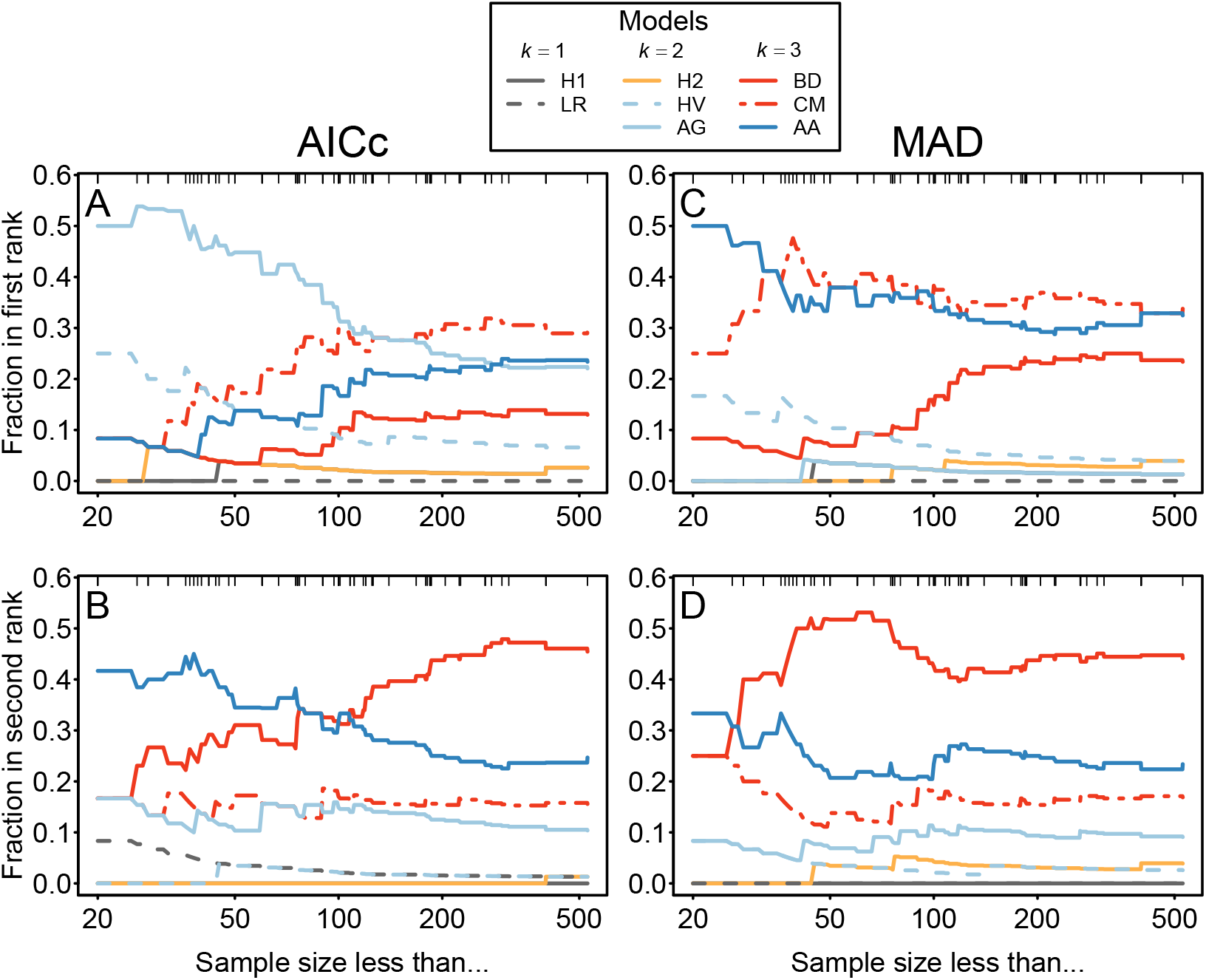
The effect of sample size on the proportion of datasets for which each of the considered functional-response models was ranked as (A) first or (B) second as judged by *AIC*_*c*_, and as (C) first or (D) second as judged by *MAD*. In each panel, more and more datasets are included in the pool of considered datasets as the maximum considered sample size increases from left to right, starting with all datasets having sample sizes of 20 or less. Tick-marks reflect the distribution of sample sizes greater than 20. Note that a model’s rank is not equivalent to the support for that model because lower-ranked models may have equivalent support (i.e. Δ*AIC*_*c*_ < 2). See Table 1 for model abbreviations.

As judged by *MAD*, the Arditi-Akçakaya model ranked first for 28 (36.4%) of all considered datasets (Fig. 2B, Table 2). This was followed by the Crowley-Martin and Beddington-DeAngelis models which respectively ranked first for 24 (31.2%) and 18 (23.2%) datasets. The Beddington-DeAngelis and Arditi-Akçakaya models were the most frequent models to be ranked second at a respective 34 (44.2%) and 18 (23.4%) times. According to *MAD*, no datasets ranked the linear ratio-dependent model as first and one ranked the Arditi-Ginzburg model as first. Forty-six (59.7%) datasets had at least one model producing a *MAD* within 1% of the first-ranked model. Of the 28 datasets for which the Arditi-Akçakaya model was ranked first, the Beddington-DeAngelis, Crowley-Martin, and Holling Type II models produced *MAD* values within 1% a respective 17 (60.7%), 8 (28.6%), and 8 (28.6%) times. The Arditi-Ginzburg, Hassell-Varley, and linear ratio-dependent models did so a respective 7 (25.0%), 2 (7.1%), and 1 (3.6%) times. The *MAD* of more complex models was not always less than that of their corresponding simpler models, in contrast to their likelihood (underlying *AIC*_*c*_) which must improve with complexity.

For *MAD*, the smaller their sample size, the greater was the proportion of datasets that ranked the Arditi-Akçakay or Hassell-Varley models as first (Fig. 3C), and the Arditi-Akçakay model as second (Fig. 3D). Correspondingly, the more that datasets with larger sample sizes were considered, the greater was the proportion of datasets that ranked either the Crowley-Martin or Beddington-DeAngelis models as first, and the Beddington-DeAngelis model as second. Datasets with larger sample sizes were no better in their ability to discriminate among models than were smaller datasets, with 16 (41.0%) of the 39 smaller datasets and 15 (39.5%) of the 38 larger datasets having a single model outperform all others by the 1% criterion (Fig. 2B).

### Parameter inference

Magnitudes of consumer interference as estimated assuming the Arditi-Akçakaya model ranged from *m* ≈ 0 to 3.8 with an overall mean of 0.94 (± 0.62 std. dev.). However, studies with smaller sample sizes generated larger interference estimates (Fig. 4) such that, for example, the mean estimate of studies with a sample size exceeding the median sample size was 0.71 (± 0.44 std. dev.) with a range of *m* ≈ 0 to 1.93. No other covariate was similarly associated with the estimated strength of consumer interference (Fig. S1). Although uncertainty around point estimates also tended to increase the lower the sample size, this increase occurred at a slower rate than did that of the point estimate magnitudes; most large-magnitude estimates associated with small sample sizes were therefore well constrained. Use of the Arditi & Akçakaya (1990) method of interference strength estimation for the subset of datasets to which it could be applied did not alter our qualitative inference regarding the presence of bias among low sample size studies (Figs. S4-S5).

**Figure 4:**
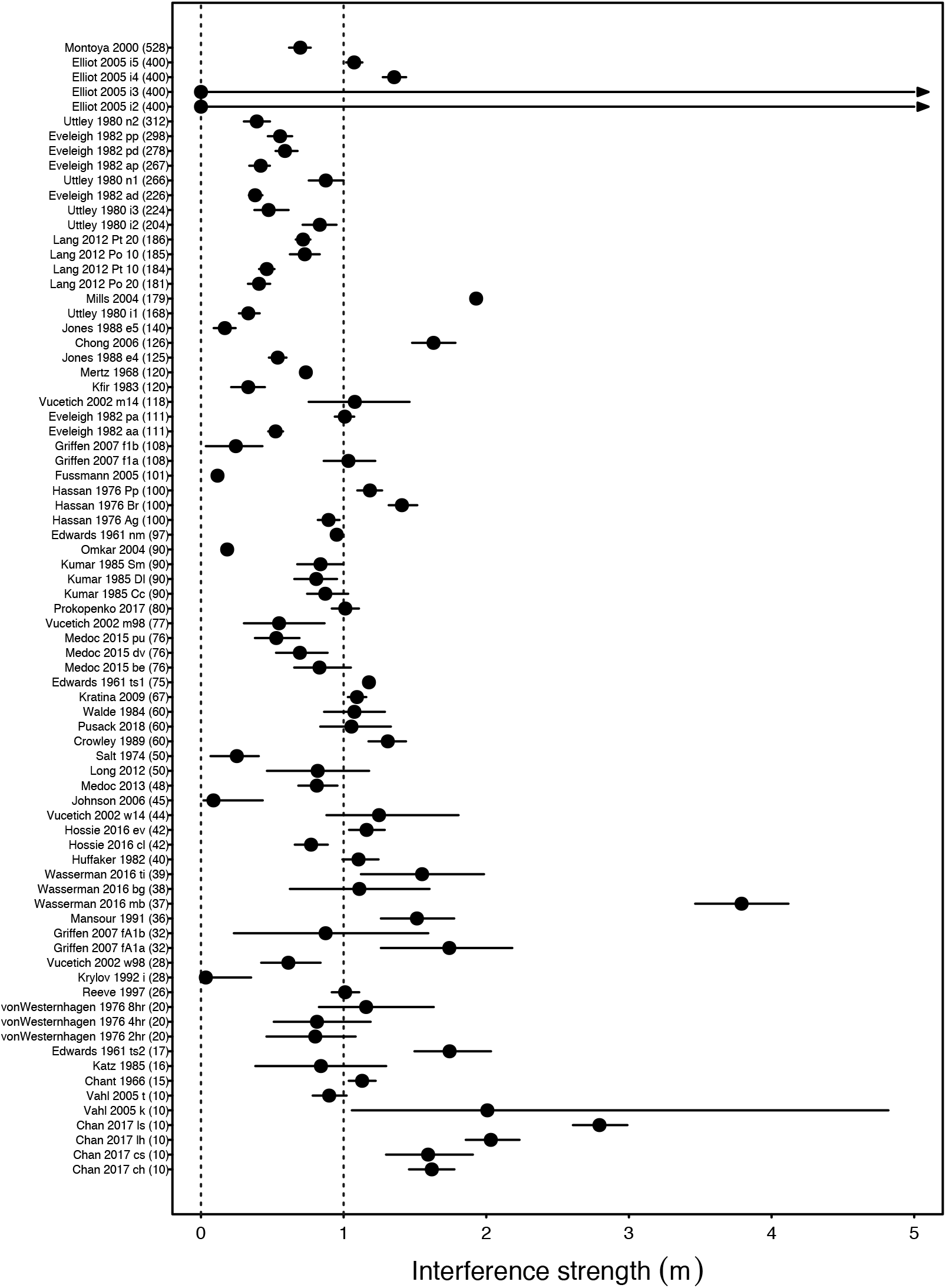
The strength of consumer interference, *m*, as estimated for each dataset assuming the Arditi-Akçakaya model, with datasets ordered by their sample size and intervals representing one standard error. The large interval for some datasets is due to poor parameter identifiability despite large sample size. See Fig. S1 for an equivalent figure on which covariate information is overlaid. Note that, due to the myriad sources of variation that exist across datasets –including differences in experimental design, observer error, and species’ biology (including the ‘true’ strength of interference) –the apparent pattern between estimates and sample sizes should not be interpreted as reflecting an underlying quantitative relationship of use for correcting parameter-estimation bias or inferring an asymptotic or mean value of interference strength.

## Discussion

Past appeals for a data-driven resolution to the debate over consumer dependence have been answered by a doubling of the number of published datasets since the clarion call of Abrams & Ginzburg (2000). Barring an exceptionally strong publication bias (Abrams, 2015; Osenberg *et al.*, 1999), our analyses of the accumulated data substantiate the pragmatic resolution of the debate’s primary focus: consumer dependence is common, and neither exact prey-nor ratio-dependent models reflect statistically-parsimonious descriptors (as judged by *AIC*_*c*_ and *MAD*) of the density dependence exhibited by most consumer feeding rates. Our analyses also confirm prior syntheses of prey-only experiments, showing that consumer feeding rates are almost invariably a saturating function of prey densities (Jeschke *et al.*, 2004).

Our primary new insight, however, is the existence of strong statistical bias in the ability of the available data to help characterize and quantify consumer dependence. First, inadequate sample sizes bias the standard information-theoretic comparison of model performance towards the models of Hassell & Varley (1969), Arditi & Ginzburg (1989), and, by extension, their generalized combination by Arditi & Akçakaya (1990). Our results on *AIC*_*c*_, paired with those on mean absolute deviation, are consistent with this bias being due to a combination of the (mis-)estimated parametric and geometric complexity that these models entail (Box 1). This bias added to the poor identifiability of *m* at small sample sizes which caused *AIC*_*c*_ to (correctly) give less support to the Arditi & Akçakaya (1990) model than the model of Arditi & Ginzburg (1989) for which *m* = 1. Second, the estimation of interference strengths by application of the long-established method of maximum likelihood to the model of Arditi & Akçakaya (1990) is biased by inadequate sample sizes towards larger-than-true values of interference strength. Although the magnitude of this estimation bias will depend on the details of each dataset, its qualitative direction is to have been anticipated (Box 2).

Both forms of bias are *not* specific to any of the functional-response models we considered, including the Arditi-Akçakaya model and its parameters. Indeed, both forms of bias may occur for all nonlinear functional-response models – and even for linear models when eaten prey are not replaced (Box 2) – regardless of whether they are consumer dependent or even only resource dependent. Unfortunately, their universality and prevalence have not been recognized in the functional-response literature, nor in the larger literatures on the characterization and estimation of species interaction strengths more generally (Wootton & Emmerson, 2005). Therefore, in what follows, we first discuss the general nature of biases that arise at low sample sizes, and the difficulty of accounting for them. Then, focusing on the results of the largest of datasets, we outline several inferences with which to guide the future development of functional-response theory and experiments. The issues we highlight emphasize the need for a more explicit consideration of the varied purposes that motivate the empirical study of consumer feeding rates and their functional forms.

### Bias is general

Although the sample size dependent bias of information criteria, like *AIC*_*c*_, and of maximum-likelihood parameter estimators, were recognized for nonlinear models long ago (Box 1 and Box 2), existing methods seeking to adjust for bias have not been adopted in the functional-response literature. The likely reason for this is that – to our knowledge – simple, off-the-shelf methods do not exist. Bias quantifica-tion remains an active area of statistical research even for linear models (Kosmidis, 2014), with the most general methods, such as bootstrapping and cross-validation (Efron & Tibshirani, 1994; Myung, 2000), requiring sample sizes greater than those used in many functional-response experiments.

The root cause of the difficulty is that bias is dependent not only upon sample size alone, but also on the combination of a given model’s functional form and the variation inherent within each given dataset. More specifically, both model-comparison bias (Box 1) and parameter-estimation bias (Box 2) are a function of a focal model’s parametric and geometric complexity, how well these are estimated, and for consumer functional responses, how these interact with the feeding rate variation within and among the treatment levels of an experiment. Sources of variation will include experimental design (e.g., how many and how large a range of species abundance levels are employed; Sarnelle & Wilson (2008); Uszko *et al.* (2020)), observer error in estimating species densities and the number of eaten prey (Jost & Arditi, 2000), and the behavioural variation that exists among consumer and prey individuals (Abrams, 2010; Chesson, 1984). Many such sources of variation will be present even in the well-controlled laboratory studies that dominate the literature, and could even be magnified in such studies given the small numbers of individual animals that are typically involved at low-abundance treatment levels (Coblentz & DeLong, 2020; Novak *et al.*, 2017).

Both forms of bias therefore pertain to functional-response models in general, including those that do not consider consumer dependence. For example, although we focused our assessment on the interference parameter *m* of the Arditi-Akçakaya model because it is dimensionless and thus more easily compared across studies, exponents like it are expected to be particularly sensitive to estimation bias because their maximum-likelihood estimators involve logarithmic functions of the data (see Box 2 and the Supplementary Materials S7). Bias is similarly probable and strong among published estimates of body mass scaling exponents and the Hill exponent in the respective contexts of allometric scaling theory and the density-dependent per capita attack rates of Type-III functional responses (e.g., Baudrot *et al.*, 2016; Hossie & Murray, 2016; Rall *et al.*, 2012). Few datasets in our compilation evidenced over-prediction at low prey densities by the models we considered (which could indicate density-dependent attack rates) and too few included information on consumer and prey body masses, hence the extent of the issue in these contexts deserves further investigation.

Importantly, parameter estimation bias will inflict all other nonlinear functional-response parameters as well. This will occur directly via the bias inherent in their own maximum-likelihood estimators (Box 2), and indirectly through their functional relationships and data-dependent covariances with the estimates of other biased parameters. For example, bias is also expected for the attack rate parameter with which the interference parameter is functionally intertwined in the Arditi-Akçakaya model (see also Gutenkunst *et al.*, 2007; Uszko *et al.*, 2020), or for any deterministic functional-response parameter when the assumed likelihood model entails “nuisance” parameters whose estimates are biased (e.g., the standard deviation of the Gaussian likelihood; see Supplementary Materials S6). Even for the simple Holling Type II model, estimates of the attack rate and handling time parameters can exhibit significant covariation (Papanikolaou *et al.*, 2016; Uszko *et al.*, 2020). The effect of covariation on bias will be especially large when feeding rate variation is large and heteroskedastic, or when the range over which species abundances are varied is insufficient to constrain estimates. The lower the sample size, the more challenging such issues of parameter identifiability become. This issue is similar to the bias caused by presuming the absence of (co)variation among parameters, as can occur when one or more parameters is fixed to a specific value (Hossie & Murray, 2016).

### Fitting models to data serves varied purposes

What can be said about consumer dependence from the datasets in our compilation that do have large sample sizes? We draw three main conclusions that are applicable to the functional-response literature more generally.

First, as judged by *AIC*_*c*_, each of the three most complex models that we considered had statistically clear support for being the sole best-performing model for several datasets, all other models having Δ*AIC*_*c*_ > 2. Therefore, consumer dependence is pervasive but there is no universally best functional form with which to characterize it (Skalski & Gilliam, 2001). Second, the strength of consumer interference as judged by fitting the Arditi-Akçakaya model varied from being very weak to very strong across consumer species and even for different life stages of the same species (e.g., Elliott, 2005). Focusing on the general tendency of *m* after establishing that consumer dependence is prevalent is therefore asking the wrong question, with future work more fruitfully considering the potential predictors of variation in mutual predator effects (DeLong, 2014; Novak *et al.*, 2017). Together, these two conclusions indicate ample room for more complex models to provide useful biological insight and to statistically outperform the models considered here (Abrams, 2015; Evans *et al.*, 2013; Stouffer & Novak, 2020). Rigorous evidence of consumers for which conspecific density dependence is entirely absent would also be useful, with much work remaining to justify the relevance of species-isolating laboratory experiments to the functional responses exhibited by consumers in nature (Novak *et al.*, 2017; Preston *et al.*, 2018).

These first two conclusions do not, however, imply that more complex functional-response models will necessarily be more useful than simpler models. Many have advocated for simple models on the basis of their analytical tractability, elegance, and utility in management contexts (e.g., Adkison, 2009; Arditi & Ginzburg, 2012; Odenbaugh, 2005). However, the third conclusion we draw from our analyses of large sample size datasets is that multiple models of varying complexity perform near equally well when performance is judged by the high but otherwise arbitrary standard of describing the data with a *MAD <* 1% of the top-performing model. Although our assessment did not evaluate the goodness-of-fit of the models in an absolute sense, for a number of datasets these “equally well-describing” models were not among the *AIC*_*c*_-judged top-performing models.

The ostensible contrast between our third and first two conclusions points to the varied purposes that motivate the fitting of functional-response models to functional-response data. Most functional-response studies have been justified by a generic desire to better describe and understand the functional form of consumer feeding rates to better predict population dynamics. Old and rejuvenated literatures on the varied and overlapping meanings of description, understanding, and prediction in ecology show-case the many ways in which this expressed desire is too simplistic (Doak *et al.*, 2008; Elliott-Graves, 2019; Evans *et al.*, 2013; Levins, 1966; Maris *et al.*, 2018; Odenbaugh, 2005; Pennekamp *et al.*, 2019; Shmueli, 2010). Although a synthesis of these terms for ecology is far from complete, pertinent highlights of the literature include evidence that simple, non-mechanistic population models can often better forecast population dynamics than even the models with which the dynamics were simulated in the first place (Perretti *et al.*, 2013); that parameter estimates need not all be well-constrained to make accurate predictions in complex systems (Gutenkunst *et al.*, 2007); that consumers may often experience a small enough range in prey population sizes that their functional responses are effectively linear under field conditions (Novak, 2010; Preston *et al.*, 2018); and that functional nonlinearities important to describing variation at some spatial, temporal, or biological scales need not be important – or indeed logical – at other scales (Chesson, 2009; Morozov & Petrovskii, 2013). Others have distinguished among various types of prediction – including explanatory, out-of-sample, extrapolatory, transferable, and forecasting forms (e.g., Maris *et al.*, 2018; Odenbaugh, 2005; Shmueli, 2010) – revealing how judgments of model performance in regards to predictive utility are by their nature dependent on choices of prediction type. Alternative motivations and choices similarly underlie vibrant discussions regarding the uses of alternative information criteria in statistical model selection (Aho *et al.*, 2014; Myung, 2000; Shmueli, 2010). Indeed, our choice to present model comparisons using *AIC*_*c*_ was motivated only by the fact that its antecedent, *AIC*, is the most commonly used information criterion in ecology (Aho *et al.*, 2014); our inferences regarding model-comparison bias at small sample sizes remain qualitatively unchanged when using *AIC* or *BIC* (Supplementary Materials S4).

In fact, the varied purposes that motivate the fitting of functional-response models to data are also relevant in regards to whether the statistical bias of low sample size studies is to be considered problematic or not. Efforts to avoid model-comparison bias through the use of complexity-penalizing information-theoretic criteria are typically driven by a desire to avoid over-fit models whose within-sample deviations may be low but whose out-of-sample prediction error is high relative to that of less complex models (Myung, 2000). However, parameterizations of a given model using biased estimators can result in lower prediction error than when unbiased estimators are used (Shmueli, 2010). This is because a model’s prediction error reflects both its bias and the variation in the data it cannot explain (i.e. a model’s accuracy and precision, respectively; Bolker, 2008), among which there can exist a trade-off. For example, even an unbiased estimator can grossly underestimate 99% of the data as long as it overestimates the remaining data even more egregiously, outliers notwithstanding. In this sense, the parameter estimation bias that is present among low sample size studies need not be considered problematic if the sole motivation is to be descriptive of a predator’s feeding behaviour in a within-sample explanatory sense.

### Conclusions

Many forms and sources of bias exist in the functional-response literature. These include recently recognized sources related to an experiment’s design and venue (Li *et al.*, 2018; Novak *et al.*, 2017; Preston *et al.*, 2018; Uiterwaal *et al.*, 2018; Uiterwaal & DeLong, 2018), as well as persistent model-associated sources (Barraquand, 2014; Damgaard, 2020; Hossie & Murray, 2016; Jost & Arditi, 2000; Marshal & Boutin, 1999; McCoy *et al.*, 2012; Morozov & Petrovskii, 2013; Novak, 2010; Novak & Wootton, 2010; Pascual & Kareiva, 1996; Rosenbaum & Rall, 2018; Trexler *et al.*, 1988). Our analyses add to this list of issues, exposing the pervasive effect that small sample sizes have had on inferences of consumer dependence, but they are also not immune to them. For example, we did not address issues related to failing to include the “true” model among the models we considered (Aho *et al.*, 2014; Burnham & Anderson, 2002). Hence, for example, our estimates of *m* assuming the Arditi & Akçakaya (1990) model may have been further biased for datasets where Type III-like density dependence was present (Hossie & Murray, 2016).

We acknowledge that our analyses also do not address the quantitative magnitude by which small sample sizes could bias model comparisons and parameter estimates. Instead we have relied on qualitative inferences alone. This is because bias correction is nontrivial and because the sources and forms of (co)variation among datasets and models are so varied that no single empirical or simulated dataset is likely to be sufficiently representative and hence universally useful. We note, however, that even seemingly small amounts of bias could correspond to sizeable biological consequences. All else being equal, overestimating the true value of *m* by a multiplicative factor of *E* is equivalent to mis-counting the number of predators by a multiplicative factor of *P*^*є*−1^. Hence, in the case of a bias as small as 5% (*є* = 1.05), the inferred interference rate between 10 predator individuals would be nearly equivalent to having had 11 individuals instead, 26 individuals would be nearly equivalent to 31, 100 individuals would be nearly equivalent to 126, and 150 individuals would be nearly equivalent to 193. While such consequences of overestimating *m* may thus seem like less of a problem for studies entailing low numbers of predators (e.g., 1, 2, or 3 predators individuals, as most studies do), these same studies also tend to be those with the lowest sample sizes. As a result, they will also tend to experience the greatest estimation bias and will tend to affect the greatest mis-predictions when extrapolating to the population-level effects of predation in nature.

For overcoming model-comparison and parameter-estimation bias there is but one simple solution: future studies must increase their sample sizes. We therefore urge authors, reviewers, and editors to hold studies to a higher standard. That said, we offer no rule of thumb for determining what sample size is enough; as noted above, the sample size necessary to effectively reduce bias will depend on the variance-generating processes of the system in question and the experimental designs and models with which these processes are studied. Continued developments in the areas of symbolic regression and optimal experimental design should have much insight to offer in reducing the logistical burden that will be required (Martin *et al.*, 2018; Moffat *et al.*, 2020; Okuyama & Bolker, 2012; Zhang *et al.*, 2018). This notwithstanding, the most fundamental advance that is needed is a conceptual one regarding a clarity of purpose. Using statistics to bridge functional-response models with empirical data requires recognizing that data, theory, and models (both deterministic and statistical) each represent necessarily incomplete and sometimes incompatibly-mismatched characterizations of nature. Hence, for some purposes, failure to fit data need not invalidate a model’s utility, while for others a model’s superior fit need not substantiate it. Clarity regarding the purpose of functional-response studies will aid in determining the map between biological objectives and methodological approach, and is needed for the functional-response concept to reliably serve as the foundation for work across population, community, and evolutionary ecology as it is often considered to be.

## Acknowledgments

We thank Stella Uiterwaal and Gregor Kalinkat for providing their compiled bibliographies of functional-response studies. For generously providing data, we thank Roger Arditi, Sven Bacher, Shane Blowes, Kevin Chan, Juang-Horng Chong, Will Cresswell, Malcolm Elliot, Gregor Fussmann, Fatemeh Ganjisaf-far, Mark Hebblewhite, Tom Hossie, Pavel Kratina, Birgit Lang, Chris Long, Anders Nilsson, Christina Prokopenko, Timothy Pusack, John Reeve, Thierry Spataro, Adrian Stier, John Vucetich, and Will White, as well as all authors who made their data available in online repositories. We further thank Peter Abrams, Roger Arditi, Lev Ginzburg, Tom Hossie, the Theoretical Breakfast group at OSU, and two anonymous reviewers for suggestions that improved the manuscript. DBS acknowledges the support of the Marsden Fund Council, the New Zealand Government (grant 16-UOC-008), and a University of Canterbury Erskine grant.

## Author contributions

Both authors contributed equally to this work. MN wrote the first draft.

## Code and data availability

Code for all analyses, as well as most datasets, are available at *https://github.com/stoufferlab/general-functional-responses*. These and additional data sets have also been posted to online repositories per agreement with data contributors, or were obtained from repositories to which they had previously been posted by the original authors (see Table S1).

## Notes

### Competing Interest Statement

The authors have declared no competing interest.

