## Supplementary Materials for "Systematic bias in studies of consumer functional responses"

### *Supplementary Materials*:
### Systematic bias in studies of consumer functional responses

Mark Novak & Daniel B. Stouffer

#### Contents

**S1 Covariates figure & summary table of datasets** **2**

**S2 Summary of prior syntheses** **6**

**S3 Model flexibility as judged by the Fisher Information Approximation** **6**

**S4 Model performance as judged by AIC & BIC** **7**

**S5 Arditi and Akçakaya method for estimating interference strengths** **11**

**S6 Dispersion and nuisance parameters as sources of bias** **13**

**S7 Maximum likelihood estimators** **13**

#### S1 Covariates figure & summary table of datasets

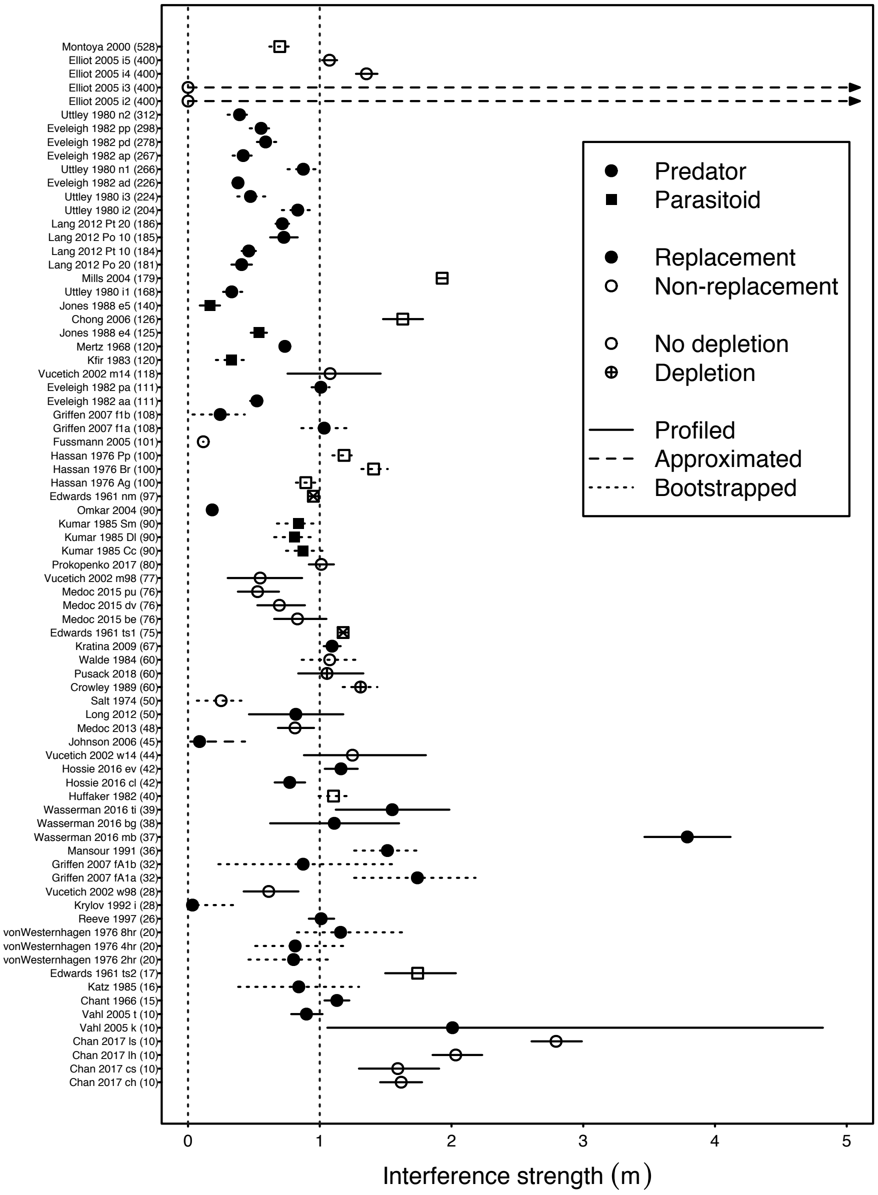

Figure S1: The estimated strength of consumer interference assuming the Arditi-Akçakaya model, with datasets ordered by their sample size and differentiated by covariate information. Non-replacement studies are designated as incurring significant prey depletion when more than 10% of replicates had more than 90% of available prey consumed. We infer that consumer type, prey depletion, and replacement scenario are not confounded with sample size.

Table S1: A summary of used datasets. "Dataset" refers to the specific experiment from the study, and '-' implies there was only one dataset available. "Raw" refers to whether we were able to use the raw data at the level of each treatment replicate, or whether we instead used means and associated uncertainty intervals to produce bootstrapped datasets. "Type" refers to whether the data was provided to us by the author, was obtained from an online repository, or was extracted from the publication. "Source" refers to the figures and tables from which the data where extracted. "Nobs" indicates the sample size. "Replaced" refers to the whether consumed prey were replaced during the study (or whether the parasitoid was considered discriminatory or not), which dictated our use of a binomial versus a Poisson likelihood.

| Study | Dataset | Raw | Type | Source | Citation | Nobs | Replaced | Consumer |
|---|---|---|---|---|---|---|---|---|
| Chan *et al.* (2017) | ch | Yes | Provided | - | Chan *et al.* (2017) | 10 | Yes | Predator |
| Chan *et al.* (2017) | cs | Yes | Provided | - | Chan *et al.* (2017) | 10 | Yes | Predator |
| Chan *et al.* (2017) | lh | Yes | Provided | - | Chan *et al.* (2017) | 10 | Yes | Predator |
| Chan *et al.* (2017) | ls | Yes | Provided | - | Chan *et al.* (2017) | 10 | Yes | Predator |
| Chant & Turnbull (1966) | - | Yes | Extracted | Table 2 | Novak & Stouffer (2020) | 15 | No | Predator |
| Chong & Oetting (2006) | - | Yes | Provided | - | Chong (2020) | 126 | Yes | Parasitoid |
| Crowley & Martin (1989) | - | No | Extracted | Fig. 2 | Novak & Stouffer (2020) | 60 | Yes | Predator |
| Edwards (1961) | nm | Yes | Extracted | Tables 1, 2 & 3 | Novak & Stouffer (2020) | 97 | Yes | Parasitoid |
| Edwards (1961) | ts1 | Yes | Extracted | Tables 1, 2 & 3 | Novak & Stouffer (2020) | 75 | Yes | Parasitoid |
| Edwards (1961) | ts2 | Yes | Extracted | Tables 1, 2 & 3 | Novak & Stouffer (2020) | 17 | Yes | Parasitoid |
| Elliott (2005) | i2 | Yes | Provided | - | Elliott (2020) | 400 | Yes | Predator |
| Elliott (2005) | i3 | Yes | Provided | - | Elliott (2020) | 400 | Yes | Predator |
| Elliott (2005) | i4 | Yes | Provided | - | Elliott (2020) | 400 | Yes | Predator |
| Elliott (2005) | i5 | Yes | Provided | - | Elliott (2020) | 400 | Yes | Predator |
| Eveleigh & Chant (1982) | aa | No | Extracted | Tables 4, 5 & 8 | Novak & Stouffer (2020) | 111 | No | Predator |
| Eveleigh & Chant (1982) | ad | No | Extracted | Tables 4, 5 & 8 | Novak & Stouffer (2020) | 226 | No | Predator |
| Eveleigh & Chant (1982) | ap | No | Extracted | Tables 4, 5 & 8 | Novak & Stouffer (2020) | 267 | No | Predator |
| Eveleigh & Chant (1982) | pa | No | Extracted | Tables 4, 5 & 8 | Novak & Stouffer (2020) | 111 | No | Predator |
| Eveleigh & Chant (1982) | pd | No | Extracted | Tables 4, 5 & 8 | Novak & Stouffer (2020) | 278 | No | Predator |
| Eveleigh & Chant (1982) | pp | No | Extracted | Tables 4, 5 & 8 | Novak & Stouffer (2020) | 298 | No | Predator |
| Fussmann *et al.* (2005) | - | Yes | Provided | - | Fussmann (2020) | 101 | Yes | Predator |
| Griffen & Delaney (2007) | f1a | No | Extracted | Fig 1 & Fig A1 | Novak & Stouffer (2020) | 108 | No | Predator |
| Griffen & Delaney (2007) | f1b | No | Extracted | Fig 1 & Fig A1 | Novak & Stouffer (2020) | 108 | No | Predator |
| Griffen & Delaney (2007) | fA1a | No | Extracted | Fig 1 & Fig A1 | Novak & Stouffer (2020) | 32 | No | Predator |
| Griffen & Delaney (2007) | fA1b | No | Extracted | Fig 1 & Fig A1 | Novak & Stouffer (2020) | 32 | No | Predator |
| Hassan (1976) | Ag | No | Extracted | Table 1 | Novak & Stouffer (2020) | 100 | Yes | Parasitoid |
| Hassan (1976) | Br | No | Extracted | Table 1 | Novak & Stouffer (2020) | 100 | Yes | Parasitoid |
| Hassan (1976) | Pp | No | Extracted | Table 1 | Novak & Stouffer (2020) | 100 | Yes | Parasitoid |

Table S1: *(continued)*

| Study | Dataset | Raw | Type | Source | Citation | Nobs | Replaced | Consumer |
|---|---|---|---|---|---|---|---|---|
| Hossie & Murray (2016) | cl | Yes | Provided | - | Hossie & Murray (2020) | 42 | No | Predator |
| Hossie & Murray (2016) | ev | Yes | Provided | - | Hossie & Murray (2020) | 42 | No | Predator |
| Huffaker & Matsumoto (1982) | - | No | Extracted | Table 1 | Novak & Stouffer (2020) | 40 | Yes | Parasitoid |
| Johnson (2006) | - | Yes | Extracted | Fig. 1 | Novak & Stouffer (2020) | 45 | No | Predator |
| Jones (1986); Jones & Hassell (1988) | e4 | Yes | Extracted | Fig. 1a & 3 | Novak & Stouffer (2020) | 125 | No | Parasitoid |
| Jones (1986); Jones & Hassell (1988) | e5 | Yes | Extracted | Fig. 1a & 3 | Novak & Stouffer (2020) | 140 | No | Parasitoid |
| Katz (1985) | - | No | Extracted | Table 1 | Arditi & Akçakaya (1990) | 112 | No | Predator |
| Kfir (1983) | - | No | Extracted | Table 1 | Novak & Stouffer (2020) | 120 | No | Parasitoid |
| Kratina *et al.* (2009) | - | Yes | Provided | - | Kratina (2020) | 67 | No | Predator |
| Krylov (1992) | i | No | Extracted | Table 1 & Fig. 1L | Novak & Stouffer (2020) | 28 | No | Predator |
| Kumar & Tripathi (1985) | Cc | No | Extracted | Tables 1 & 2 & Figs. 1 & 2 | Novak & Stouffer (2020) | 90 | No | Parasitoid |
| Kumar & Tripathi (1985) | Dl | No | Extracted | Tables 1 & 2 & Figs. 1 & 2 | Novak & Stouffer (2020) | 90 | No | Parasitoid |
| Kumar & Tripathi (1985) | Sm | No | Extracted | Tables 1 & 2 & Figs. 1 & 2 | Novak & Stouffer (2020) | 90 | No | Parasitoid |
| Lang *et al.* (2012) | Po 10 | Yes | Provided | - | Lang (2020) | 185 | No | Predator |
| Lang *et al.* (2012) | Po 20 | Yes | Provided | - | Lang (2020) | 181 | No | Predator |
| Lang *et al.* (2012) | Pt 10 | Yes | Provided | - | Lang (2020) | 184 | No | Predator |
| Lang *et al.* (2012) | Pt 20 | Yes | Provided | - | Lang (2020) | 186 | No | Predator |
| Long *et al.* (2012) | - | Yes | Provided | - | Long (2020) | 50 | No | Predator |
| Mansour & Lipcius (1991) | - | No | Extracted | Fig. 1a | Novak & Stouffer (2020) | 36 | No | Predator |
| Médoc *et al.* (2013) | - | Yes | Provided | - | Médoc *et al.* (2020b) | 48 | Yes | Predator |
| Médoc *et al.* (2015) | be | Yes | Provided | - | Médoc *et al.* (2020a) | 76 | Yes | Predator |
| Médoc *et al.* (2015) | dv | Yes | Provided | - | Médoc *et al.* (2020a) | 76 | Yes | Predator |
| Médoc *et al.* (2015) | pu | Yes | Provided | - | Médoc *et al.* (2020a) | 76 | Yes | Predator |
| Mertz & Davies (1968) | - | Yes | Extracted | Table 1 | Novak & Stouffer (2020) | 120 | No | Predator |
| Mills & Lacan (2004) | - | Yes | Extracted | Fig. 1a-c | Novak & Stouffer (2020) | 179 | Yes | Parasitoid |
| Montoya *et al.* (2000) | - | No | Extracted | Table 1 | Novak & Stouffer (2020) | 528 | Yes | Parasitoid |
| Omkar & Pervez (2004) | - | Yes | Provided | - | Omkar & Pervez (2004) | 90 | No | Predator |
| Prokopenko *et al.* (2017) | - | Yes | Provided | - | Prokopenko (2020) | 80 | Yes | Predator |
| Pusack *et al.* (2018) | - | Yes | Provided | - | Pusack (2020) | 60 | Yes | Predator |
| Reeve (1997) | - | Yes | Provided | - | Reeve (2020) | 26 | No | Predator |
| Salt (1974) | - | No | Extracted | Fig. 3 & Table 1 | Novak & Stouffer (2020) | 50 | Yes | Predator |
| Uttley (1980) | i1 | No | Extracted | Figs. 4.3.1, 4.3.2, 4.3.3, 8.3.1 & 8.3.2 | Novak & Stouffer (2020) | 168 | No | Predator |
| Uttley (1980) | i2 | No | Extracted | Figs. 4.3.1, 4.3.2, 4.3.3, 8.3.1 & 8.3.2 | Novak & Stouffer (2020) | 204 | No | Predator |
| Uttley (1980) | i3 | No | Extracted | Figs. 4.3.1, 4.3.2, 4.3.3, 8.3.1 & 8.3.2 | Novak & Stouffer (2020) | 224 | No | Predator |
| Uttley (1980) | n1 | No | Extracted | Figs. 4.3.1, 4.3.2, 4.3.3, 8.3.1 & 8.3.2 | Novak & Stouffer (2020) | 266 | No | Predator |
| Uttley (1980) | n2 | No | Extracted | Figs. 4.3.1, 4.3.2, 4.3.3, 8.3.1 & 8.3.2 | Novak & Stouffer (2020) | 312 | No | Predator |
| Vahl *et al.* (2005) | k | Yes | Extracted | Fig. 4 | Novak & Stouffer (2020) | 10 | No | Predator |
| Vahl *et al.* (2005) | t | Yes | Extracted | Fig. 4 | Novak & Stouffer (2020) | 10 | No | Predator |

Table S1: *(continued)*

| Study | Dataset | Raw | Type | Source | Citation | Nobs | Replaced | Consumer |
|---|---|---|---|---|---|---|---|---|
| Von Westernhagen & Rosenthal (1976) | 2hr | No | Extracted | Fig. 3 | Novak & Stouffer (2020) | 80 | No | Predator |
| Von Westernhagen & Rosenthal (1976) | 4hr | No | Extracted | Fig. 3 | Novak & Stouffer (2020) | 120 | No | Predator |
| Von Westernhagen & Rosenthal (1976) | 8hr | No | Extracted | Fig. 3 | Novak & Stouffer (2020) | 100 | No | Predator |
| Vucetich *et al.* (2002) | m14 | Yes | Provided | - | Jost *et al.* (2005); Vucetich *et al.* (2002) | 118 | Yes | Predator |
| Vucetich *et al.* (2002) | m98 | Yes | Provided | - | Jost *et al.* (2005); Vucetich *et al.* (2002) | 77 | Yes | Predator |
| Vucetich *et al.* (2002) | w14 | Yes | Provided | - | Jost *et al.* (2005); Vucetich *et al.* (2002) | 44 | Yes | Predator |
| Vucetich *et al.* (2002) | w98 | Yes | Provided | - | Jost *et al.* (2005); Vucetich *et al.* (2002) | 28 | Yes | Predator |
| Walde & Davies (1984) | - | No | Extracted | Fig 2 & 4 | Novak & Stouffer (2020) | 60 | Yes | Predator |
| Wasserman *et al.* (2016b) | bg | Yes | Repository | - | Wasserman *et al.* (2016a) | 38 | No | Predator |
| Wasserman *et al.* (2016b) | mb | Yes | Repository | - | Wasserman *et al.* (2016a) | 37 | No | Predator |
| Wasserman *et al.* (2016b) | ti | Yes | Repository | - | Wasserman *et al.* (2016a) | 39 | No | Predator |

#### S2 Summary of prior syntheses

Several notable syntheses of consumer-dependent functional-response experiments have been performed over the decades. Here we briefly summarize these syntheses to highlight their assumed statistical models and the varied sample sizes of the datasets they considered.

Hassell & Varley (1969) and Hassell (1971) analyzed 9 datasets having sample sizes ranging from $\sim$16 to 84 replicates. They provided evidence for consumer dependence by applying ordinary linear least squares regression to log-transformed 'area of discovery' (i.e. attack rate) estimates inferred from counts of the number of hosts surviving parasitism as a function of parasite abundance and initial host abundance assuming linear prey dependence and multiplicative consumer dependence.

Arditi & Akçakaya (1990) showed how the Hassell & Varley approach will underestimate interference when consumers experience non-negligible handling times or when parasites can discriminate among already parasitized and non-parasitized hosts. They compiled 15 datasets of predators and parasitoids having sample sizes ranging from 9 to 75 and used both linear and nonlinear least squares regression in estimating the multiplicative effect of consumer abundance on log-transformed attack rates, which were themselves each estimated on the basis of Holling Type II prey dependence at fixed predator abundances.

Skalski & Gilliam (2001) compiled a total of 19 datasets having sample sizes ranging from 5 to 120 (they used treatment means as data for most datasets). They evaluated the relative performance of several alternative prey-, ratio- and consumer-dependent functional-response models – including multiplicative and additive models of consumer dependence – using the bootstrap-estimated confidence intervals of parameter point estimates (for nested models differing in their number of parameters, $k$) and likelihood-ratio tests (when comparing equal-$k$ models). Their model-fitting used the method of maximum likelihood assuming log-normally distributed feeding rate residuals (i.e. minimizing squared differences between observed and predicted values, analogous to least squares regression).

Most recently, DeLong & Vasseur (2011) focused on the Arditi-Akçakaya model and combined estimates of $m$ reported in the literature (including those of prior syntheses and studies published thereafter) with estimates obtained by (re-)analyzing many datasets for a total of 51 datasets with sample sizes ranging from 9 to greater than 179. In estimating interference strength, they applied four methods – including those of Hassell & Varley (1969) and Arditi & Ginzburg (1989) – using ordinary least squares regression and either log-transformed feeding rate or attack rate estimates as the response variable. (Their analysis also included 9 datasets entailing metabolic rate as the response variable.)

#### S3 Model flexibility as judged by the Fisher Information Approximation

We consider model flexibility through the lens of the Minimum Description Length principle as implemented via the Fisher Information Approximation (*FIA*). We refer readers to Myung *et al.* (2006), Ly *et al.* (2017) and references therein for more extensive summaries and further details.

Derived by Rissanen (1996), *FIA* is one of many information criterion similar to *AIC* and $AIC_c$ that penalize the likelihood of a fitted statistical model by its complexity. While *AIC* and $AIC_c$ do so purely on the basis of the model's $k$ free parameters and the sample size $n$ of the data (for the latter), *FIA* does so by also considering the model's mathematical flexibility with respect to the structure of the data. That is,

$$\begin{aligned} AIC = & \quad -2\ln\mathcal{L}(\theta|y) \quad + \ 2k & \text{(S1)} \\ AIC_c = & \quad -2\ln\mathcal{L}(\theta|y) \quad + \ 2k + \frac{2k(k+1)}{n-k-1} & \text{(S2)} \\ FIA = & \quad -\ln\mathcal{L}(\theta|y) \quad + \frac{k}{2}\ln\left(\frac{n}{2\pi}\right) \ + \ \ln\int_\theta \sqrt{\det I(\theta)}\,\mathrm{d}\theta. & \text{(S3)} \end{aligned}$$

The first term of each criterion is the negative log-likelihood of the model (with parameter(s) $\theta$) given the data $y$. It reflects a measure of the model's *goodness-of-fit* to the data. The second term of each criterion reflects the model's *parametric complexity* (a.k.a. its dimensionality), which is independent of the data beyond the influence of $n$. The third term of *AIC*$_c$ is a correction factor for *AIC*'s mis-estimation of parametric complexity at low sample sizes.

The third term of *FIA* reflects the model's *flexibility* (a.k.a. its geometric or structural complexity). The Expected Fisher information matrix $I(\theta)$ in the third term of *FIA* is calculated as the expectation of the Fisher information matrix, which may itself be calculated as the Hessian of the model's negative log-likelihood function (taking second-order derivatives of the function with respect to each of the model's parameters). The domain of the integral in the third term reflects the range of values that the model's parameters may exhibit; for some models this integral may not be finite and for some experimental designs the range of empirically observable parameter values may be more restricted than is mathematically conceivable.

A model's flexibility as assessed by *FIA* therefore reflects both its functional form (as encapsulated by $I(\theta)$) and the range of parameter values which are considered empirically possible. Through $I(\theta)$, flexibility is sensitive to the experimental design (i.e. treatment levels) and the identifiability of the model's parameters (i.e. how much information they share). The same two models can exhibit different relative amounts of flexibility for different experimental designs. However, different parameterizations of the same functional form (e.g., the Holling and Michaelis-Menten forms of the Type II functional response) will exhibit the same flexibility for a given experimental design when the permissible range of their parameters is equivalently limited.

#### S4 Model performance as judged by AIC & BIC

As noted in Box 1 of the main text, *AIC* (eqn. S1) has become the most frequently used information criterion for evaluating relative model performance (Aho *et al.*, 2014). It estimates the expected relative Kullback-Leibler divergence of a focal model from the true model responsible for generating the data. *AIC*$_c$ (eqn. S2) incorporates a correction for the bias that *AIC* exhibits in this estimation at small sample sizes, imposing a greater penalty on a model's parametric complexity than does *AIC*. The difference between *AIC* and *AIC*$_c$ diminishes as sample size increases.

The most commonly used alternative criterion to *AIC* is the Bayesian Information Criterion, BIC (a.k.a. Schwarz Information Criterion; Schwarz, 1978), calculated as

$$BIC = -2\ln\mathcal{L}(\theta|y) + \quad k\ln(n). \tag{S4}$$

Just as in eqns. S1-S3, the first term of eqn. S4 describes the model's goodness-of-fit per its likelihood, and the second term measures the model's parametric complexity. Unlike for *AIC* and *AIC*$_c$, but similar

Table S2: The number of datasets for which each functional response model achieved a given rank relative to all other models as judged by $AIC$ or by $BIC$.

| Rank | H1 | H2 | BD | CM | LR | HV | AG | AA |
|---|---|---|---|---|---|---|---|---|
| | | | | $AIC$ | | | | |
| 1 | 2 | 2 | 10 | 24 | 0 | 5 | 14 | 20 |
| 2 | 0 | 1 | 35 | 11 | 0 | 1 | 10 | 19 |
| 3 | 0 | 1 | 19 | 17 | 2 | 4 | 7 | 27 |
| 4 | 1 | 14 | 8 | 19 | 1 | 1 | 24 | 9 |
| 5 | 3 | 28 | 2 | 3 | 4 | 24 | 12 | 1 |
| 6 | 9 | 4 | 3 | 3 | 18 | 37 | 2 | 1 |
| 7 | 34 | 19 | 0 | 0 | 12 | 5 | 7 | 0 |
| 8 | 28 | 8 | 0 | 0 | 40 | 0 | 1 | 0 |
| | | | | $BIC$ | | | | |
| 1 | 3 | 3 | 7 | 23 | 0 | 4 | 21 | 16 |
| 2 | 0 | 1 | 34 | 11 | 1 | 2 | 8 | 20 |
| 3 | 0 | 1 | 22 | 17 | 1 | 4 | 4 | 28 |
| 4 | 0 | 14 | 8 | 20 | 1 | 1 | 23 | 10 |
| 5 | 3 | 27 | 3 | 3 | 4 | 24 | 11 | 2 |
| 6 | 10 | 3 | 3 | 3 | 18 | 37 | 2 | 1 |
| 7 | 33 | 20 | 0 | 0 | 12 | 5 | 7 | 0 |
| 8 | 28 | 8 | 0 | 0 | 40 | 0 | 1 | 0 |

to *FIA* (Section S3), a model's parametric complexity as estimated by *BIC* increases with sample size $n$. Asymptotically, *BIC* converges to twice the value of *FIA* as sample size increases (Myung, 2000).

The debate over whether to use *AIC* versus *BIC* for judging model performance runs deep among ecologists, and among statisticians more generally as well (Aho *et al.*, 2014; Vrieze, 2012). Philosophically speaking, the two criteria differ most fundamentally in their intended purpose. *AIC* is designed to select the best out-of-sample predictive model; given certain assumptions, it is asymptotically *efficient* as sample size increases (Shibata, 1983). In contrast, *BIC* is designed to select the true model; given certain assumptions, it is asymptotically *consistent* as sample size increases (Schwarz, 1978). Among their differing assumptions is that *BIC* presumes the true model to be among the considered models while *AIC* does not. Pragmatically speaking, *BIC* penalizes complex models to a greater extent than does *AIC* (when $n \geq 8$) and therefore favors models with fewer parameters.

Although these and other distinctions between the two criteria (and others) are important and directly relevant to the varied purposes that motivate the study of consumer functional responses, we do not seek to rehash nor weigh in on the debate over their use here. We do point out that although it has become common in some ecological subfields to apply both criteria and to interpret the strength of inferences accordingly, others have argued forcefully against this (Aho *et al.*, 2014). Nonetheless, for comparison purposes, and having chosen to present the results of $AIC_c$ in the main text due to its convergence on the predominantly-used criterion, *AIC*, we present the results of repeating our assessment of small sample size model-comparison bias for *AIC* and *BIC* (Table S2 and Figs. S2-S3). These generally bear out the expected favoring of simpler models by *BIC* relative to *AIC*. More importantly, they also evince the small sample size model-comparison bias that is expected for both criteria because both are only asymptotically optimal as sample size increases.

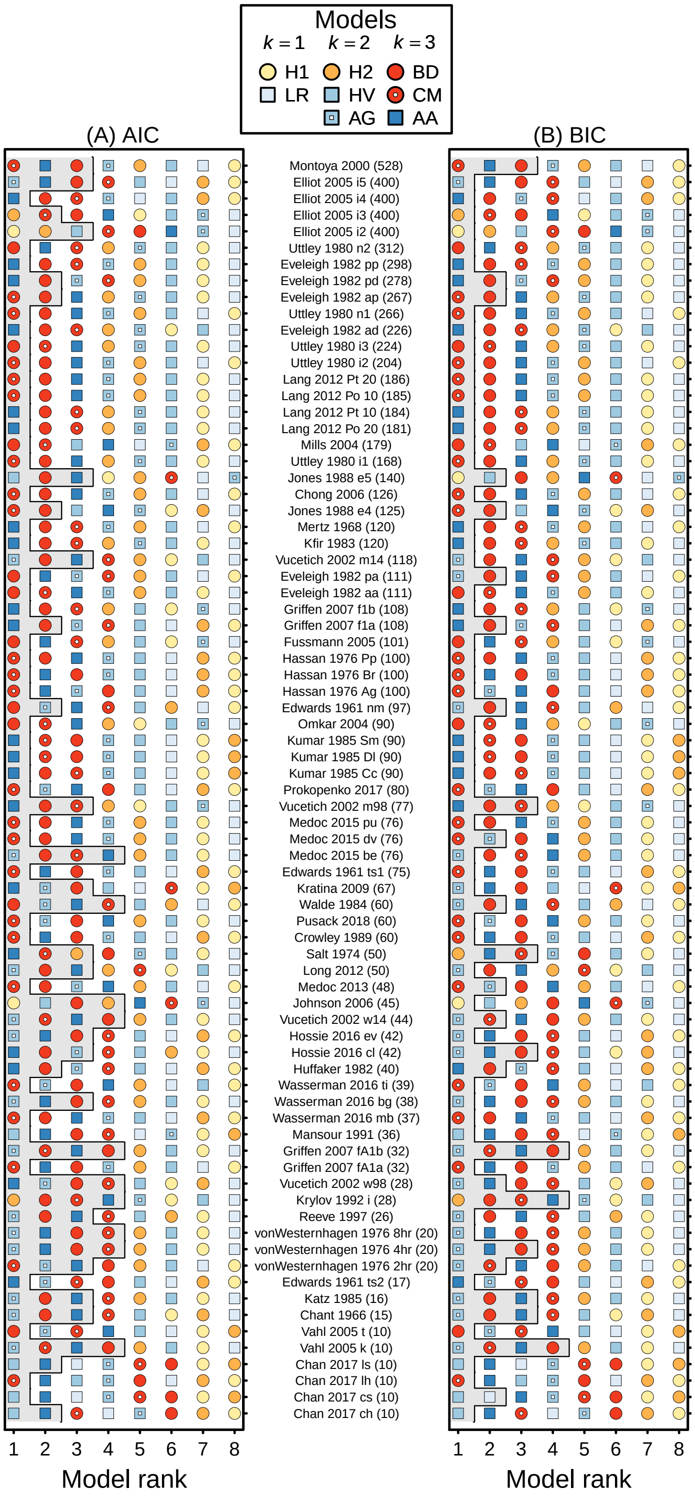

Figure S2: The rank-order performance of functional-response models as judged by (A) *AIC* and (B) *BIC*, with datasets ordered by their sample size. The gray region in (A) and (B) demarcates models interpreted as having equivalent support to the top model (i.e. $\Delta AIC < 2$ or $\Delta BIC < 2$).

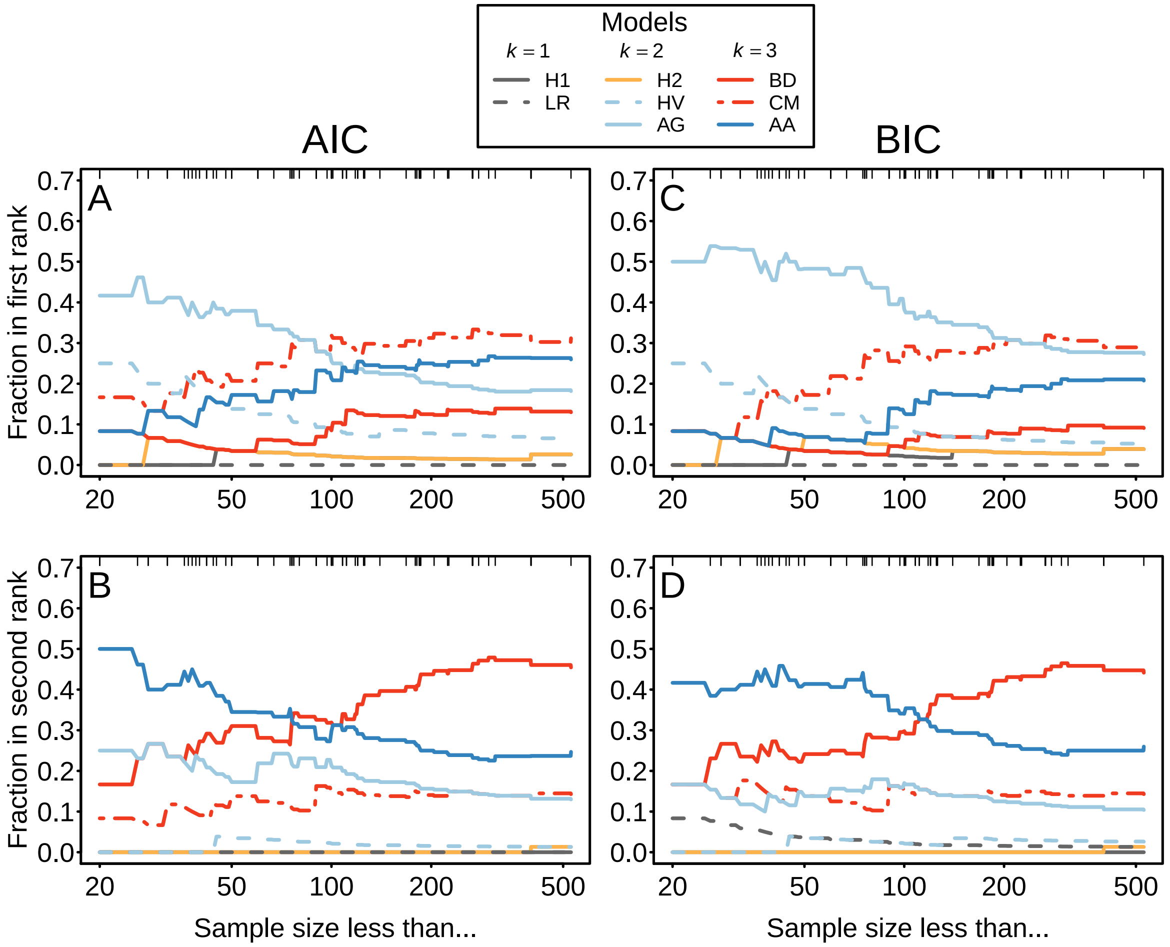

Figure S3: The effect of sample size on the proportion of datasets for which each of the considered functional-response models was ranked as (A) first or (B) second as judged by *AIC*, and as (C) first or (D) second as judged by *BIC*. In each panel, more and more datasets are included in the pool of considered datasets as the maximum considered sample size increases from left to right, starting with all datasets having sample sizes of 20 or less. Tick-marks reflect the distribution of sample sizes greater than 20. Note that a model's rank is not equivalent to the support for that model because lower-ranked models may have equivalent support.

#### S5 Arditi and Akçakaya method for estimating interference strengths

We also employed the method used by Arditi & Akçakaya (1990) to assess the degree to which past inferences using this method may have been influenced by parameter estimation bias. Although more recent studies suggest that the sequential nature of their approach is subject to additional forms of estimation bias (e.g., Uszko *et al.*, 2020), Arditi and Akçakaya used their approach ("method 2") to avoid the underestimation of interference associated with the use of the Hassell-Varley model when feeding rates saturate at high prey abundances.

The method requires discrete consumer abundance levels (treatments) with variation in prey abundances within each, and hence could only be applied to a subset of experimental datasets in our compilation. It entails the estimation of attack rates for each consumer abundance level assuming a Holling Type II model with a single handling time parameter that is common to all levels. In applying the approach, we assumed either a within-level binomial or Poisson distribution, as appropriate based on non-replacement or replacement, rather than using nonlinear least squares as done by Arditi & Akçakaya (1990). The interference parameter $m$ is subsequently estimated by the slope of a weighted linear least squares regression of log-transformed attack rates on consumer abundances, weighting each attack rate estimate by the inverse of its estimated variance. In a few cases where a given attack rate's variance could not be estimated, we used twice the value of the largest variance observed among the other attack rate estimates to weight the given attack rate's value.

As illustrated in Fig. S4, the Arditi and Akçakaya method underestimated interference strengths only slightly for most datasets (relative to the methods of our main text), and provided anomalously small estimates for some datasets. Although the number of datasets is much reduced, the sample size bias of interference estimates remains apparent (Fig. S5)

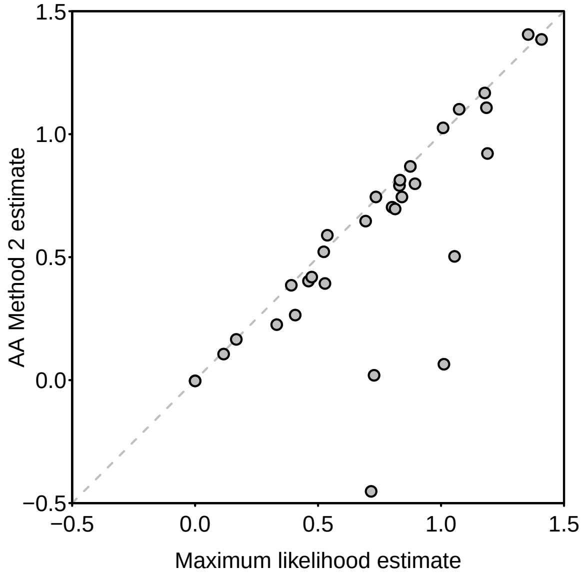

Figure S4: Interference strengths ($m$) as estimated using the methods of the main text compared to those obtained with the method used by Arditi & Akçakaya (1990) for the subset of datasets to which the latter could be applied.

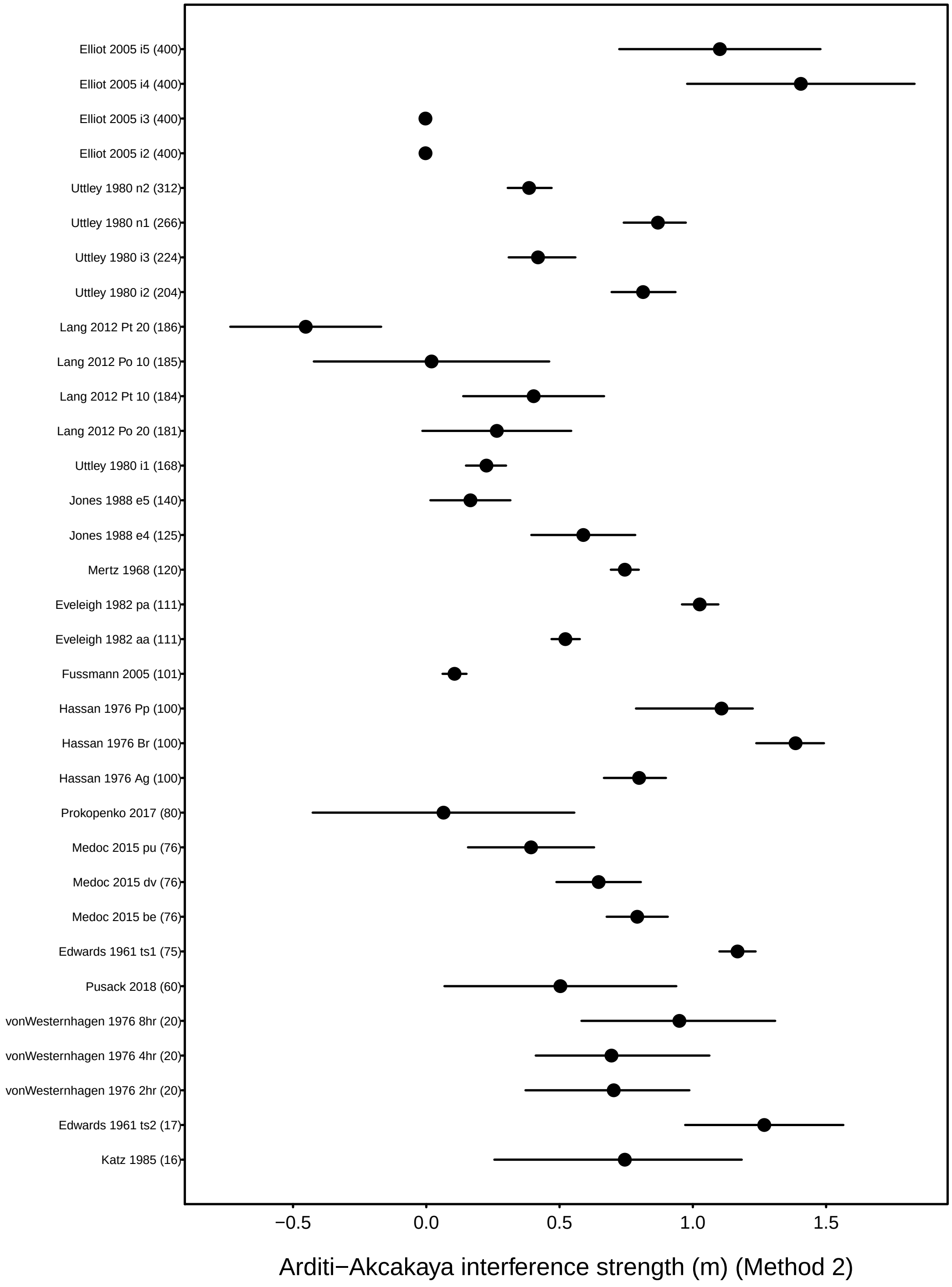

Figure S5: The strength of consumer interference as estimated with the method used by Arditi & Akçakaya (1990) for the datasets to which this method could be applied. Datasets are ordered by their sample size.

#### S6 Dispersion and nuisance parameters as sources of bias

Aside from issues of bias associated with the mathematical forms of alternative functional-response models discussed in the main text, there are also issues of bias associated with their statistical forms. More specifically, while the debate over consumer-dependent functional responses has centered on the first-principles validity and the performance of alternative deterministic models (Abrams, 2015; Arditi & Ginzburg, 2012), necessary discussions of the assumed formulation of the associated probabilistic models used in fitting models to data have been left on the periphery (Barraquand, 2014; Trexler *et al.*, 1988).

Our use of binomial and Poisson likelihood models, for example, has become standard for fitting functional responses because they more appropriately represent the generative process underlying observed counts of eaten prey at the scales at which most experiments are conducted (Bolker, 2008; Trexler *et al.*, 1988). Unlike for the Normal likelihood, the expected number of prey eaten is constrained to be positive for the Poisson and binomial, even at the low prey densities necessitated by functional-response experiments (Sarnelle & Wilson, 2008). The Poisson and binomial also do not prescribe the feeding propensities of individual consumers to be self-multiplicative as is assumed under the log-Normal likelihood, which has seen much use in prior functional-response syntheses. Rather, both the Poisson and binomial distributions become approximately normal only when the expected number of prey eaten is large.

Nevertheless, Fenlon & Faddy (2006) and others have argued for the use of likelihood models like the beta-binomial and the negative binomial as respective alternatives to the binomial and Poisson in fitting functional-response models. This is because use of binomial and Poisson likelihoods assumes that all prey individuals have an equal and temporally constant density-dependent probability of being eaten by each consumer. More specifically, the binomial and Poisson assume the variance in the data to be proportional to the mean, which is often not true for functional responses (e.g., Papanikolaou *et al.*, 2016). (It is only as the number of successes, $k$, becomes large that the variance of the negative binomial, $\mu + \mu^2/k$, approaches its mean whereby it approaches the Poisson distribution; this occurs approximately $k > 10$.) Greater-than-expected variance can bias parameter estimates when it is ignored, so use of the beta-binomial and the negative binomial is easily justified. On the other hand, the additional dispersion parameter introduced by the beta-binomial and the negative binomial to separate the variance from the mean can also bias the point estimates of the focal functional response parameters of interest when sample sizes are low, and can suck up variation in the data to artificially shrink their inferred uncertainty. This applies to the standard deviation parameter of the normal distribution as well.

Although the fitting of models using beta-binomial or negative binomial likelihoods to account for over-dispersion did not alter our primary inferences regarding the sample size dependent bias of model comparisons and parameter estimation, and are therefore omitted from our presentation, many datasets do show evidence over-dispersion. We therefore echo Fenlon & Faddy (2006) and Billiard *et al.* (2018) in encouraging future work to better characterize and utilize both the mean and variance of consumer functional responses.

#### S7 Maximum likelihood estimators

We here provide derivations of the maximum likelihood estimators (MLEs) for the parameters of a representative set of functional-response models to evidence the nature of their estimation bias. We follow the heuristic simplification of Box (1971) in assuming that independent replicate observations are made at a single treatment level having $N$ available prey and $P$ consumers. However, the qualitative nature of the bias that we infer for each of the MLEs is not specific to this assumption.

We provide derivations for each of three likelihood models, considering the counts (or proportions) of prey eaten to be random variables drawn from either a Poisson, a binomial, or a (log-)normal distribution. The Poisson applies when eaten prey are continually replaced. The binomial applies when eaten prey are not replaced (and for which $N$ reflects the initial number of available prey). The normal and log-normal are typically used when the Poisson and binomial are not applicable because feeding data are expressed as continuous rates rather than counts of prey eaten. In principal, both may be applied in situations where eaten prey are replaced and when eaten prey are not replaced.

For clarity, we first provide MLE derivations for each distribution's generic parameter(s), then apply the same procedures in deriving the MLEs for the parameters of each functional-response model. Throughout we use $y_i$ to represent the $i$th of $n$ total data points. We use $F(N, P, T, \theta)$ to represent a given functional-response model with parameter(s) $\theta$. The function $F(N, P, T, \theta)$ gives the expected number of prey eaten by $P$ consumers in the experimental time period $T$ given an (initial) abundance of $N$ prey.

#### S7.1 Poisson

The Poisson distribution has a single parameter, $\lambda$, reflecting the expected number of events as well as their variance. Given the Poisson's probability mass function

$$f(y|\lambda) = \frac{\lambda^y e^{-\lambda}}{y!}\,, \tag{S5}$$

its likelihood function is given by

$$\mathcal{L}(\lambda|y) = \prod_{i=1}^{n} f(y_i) = \prod_{i=1}^{n} \frac{\lambda^{y_i} e^{-\lambda}}{y_i!} = e^{-n\lambda} \frac{\lambda^{\sum_{i=1}^{n} y_i}}{\prod_{i=1}^{n} y_i!} \tag{S6}$$

such that its log-likelihood is given by

$$\ln \mathcal{L}(\lambda|y) = -n\lambda + \sum_{i=1}^{n} y_i \ln(\lambda) - \ln\left(\prod_{i=1}^{n} y_i!\right). \tag{S7}$$

The MLE, $\hat{\lambda}$, is obtained by finding the maximum of the log-likelihood and solving for $\lambda$:

$$\frac{d\ \ln \mathcal{L}(\lambda|y)}{d\lambda} = -n + \frac{1}{\lambda}\sum_{i=1}^{n} y_i = 0 \tag{S8}$$

$$\implies \hat{\lambda} = \frac{\sum_{i=1}^{n} y_i}{n} = \bar{y}\,. \tag{S9}$$

Hence, for the Poisson, $\hat{\lambda}$ corresponds to the arithmetic mean of the data, $\bar{y}$. We obtain the MLEs for the parameters of any given functional response by substituting $\lambda = F(N, P, T, \theta)$ into the likelihood.

***Holling Type I***

$$\lambda = F(N, P, T, \theta) = aNPT \tag{S10}$$

$$\ln \mathcal{L}(\theta|y) = -n(aNPT) + \ln(aNPT)\sum_{i=1}^{n} y_i - \ln\left(\prod_{i=1}^{n} y_i!\right) \tag{S11}$$

$$\frac{d\ \ln \mathcal{L}(\theta|y)}{da} = -nNPT + \frac{1}{a}\sum_{i=1}^{n} y_i = 0 \tag{S12}$$

$$\implies \hat{a} = \frac{\sum_{i=1}^{n} y_i}{nNPT} = \frac{\bar{y}}{NPT} \tag{S13}$$

To determine the bias in $\hat{a}$ we determine its expected value given the predicted distribution of $y_i$. If there is no bias, then $\hat{a} = a$. When the $y_i$ are Poisson random variables, checking this condition equates to evaluating

$$E(\hat{a}) = \sum_{y=0}^{\infty} \hat{a}\frac{\lambda^y e^{-\lambda}}{y!}\,. \tag{S14}$$

Note that the discrete nature of a Poisson distribution makes this a sum as opposed to an integral. Because of the linearity of expectation $E(\hat{a}) = E\left(\frac{y_i}{NPT}\right)$ and there is no need to determine the average across multiple values of $y_i$. We therefore substitute our expressions for $\hat{a}$ and $\lambda$ to obtain

$$\begin{aligned} E(\hat{a}) &= \sum_{y=0}^{\infty}\left(\frac{y}{NPT}\right)\frac{(aNPT)^y\, e^{-aNPT}}{y!} && \text{(S15)} \\ &= \frac{e^{-aNPT}}{NPT}\sum_{y=0}^{\infty}\frac{y\,(aNPT)^y}{y!} && \text{(S16)} \\ &= \frac{e^{-aNPT}}{NPT} aNPTe^{aNPT} && \text{(S17)} \\ &= a && \text{(S18)} \end{aligned}$$

and hence there is *no* bias associated with the estimator $\hat{a}$ for the Holling Type I response in this case.

The derivation of the MLE for $\alpha$ of the ratio-dependent model is equivalent and evidences that $\hat{\alpha} = \bar{y}/(NT)$, which is unbiased.

***Holling Type II***

$$\lambda = F(N,P,T,\theta) = \frac{aN}{1+ahN}PT \tag{S19}$$

$$\ln \mathcal{L}(\theta|y) = -n\frac{aNPT}{1+ahN} + \ln\frac{aNPT}{1+ahN}\sum_{i=1}^{n} y_i - \ln\left(\prod_{i=1}^{n} y_i!\right) \tag{S20}$$

$$\frac{d\ \ln \mathcal{L}(\theta|y)}{da} = \frac{\sum_{i=1}^{n} y_i + ahN\sum_{i=1}^{n} y_i - anPTN}{a(1+ahN)^2} = 0 \tag{S21}$$

$$\implies \hat{a} = \frac{\bar{y}}{N(PT - h\bar{y})} \tag{S22}$$

$$\frac{d\ \ln \mathcal{L}(\theta|y)}{dh} = 0 \implies \hat{h} = \frac{anNPT - \sum_{i=1}^{n} y_i}{aN\sum_{i=1}^{n} y_i} = \frac{aNPT - \bar{y}}{aN\bar{y}} = \frac{PT}{\bar{y}} - \frac{1}{aN} \tag{S23}$$

We check for bias in $\hat{a}$ as above with

$$E(\hat{a}) = \sum_{y=0}^{\infty} \hat{a}\frac{\lambda^y e^{-\lambda}}{y!} \tag{S24}$$

$$= \sum_{y=0}^{\infty}\left(\frac{y}{N(PT-hy)}\right)\frac{\lambda^y e^{-\lambda}}{y!}\,. \tag{S25}$$

This sum is not convergent. However, we note that Jensen's inequality for *convex* functions of $y$ dictates that

$$E\left(\frac{y}{N(PT-hy)}\right) \geq \frac{E(y)}{N(PT - hE(y))}\,. \tag{S26}$$

By substituting in $E(y) = \lambda = F(N,P,T,\theta)$ we find that

$$E(\hat{a}) \geq \frac{\lambda}{N(PT-h\lambda)} = \frac{\frac{aN}{1+ahN}PT}{N(PT - h\frac{aN}{1+ahN}PT)} = a\,, \tag{S27}$$

which means that the bias on $\hat{a}$ will always be positive.

We similarly check for bias in $\hat{h}$ with

$$E(\hat{h}) = \sum_{y=0}^{\infty} \hat{h}\frac{\lambda^y e^{-\lambda}}{y!} \tag{S28}$$

$$= \sum_{y=0}^{\infty}\left(\frac{PT}{y} - \frac{1}{aN}\right)\frac{\lambda^y e^{-\lambda}}{y!}\,, \tag{S29}$$

which is also a non-convergent sum. Jensen's inequality is again useful because $\hat{h}$ is also a *convex* function of $y$, thus

$$E\left(\frac{PT}{y} - \frac{1}{aN}\right) \geq \frac{PT}{E(y)} - \frac{1}{aN}\,. \tag{S30}$$

Substituting $E(y) = \lambda = F(N,P,T,\theta)$ into this expression we find that

$$E(\hat{h}) \geq \frac{PT}{\lambda} - \frac{1}{aN} = \frac{PT}{\frac{aN}{1+ahN}PT} - \frac{1}{aN} = h\,, \tag{S31}$$

which implies that the bias on $\hat{h}$ will always be positive.

The MLEs for the parameters of the Arditi-Ginzburg model are similar in form and bias:

$$E(\hat{\alpha}) = \quad E\left(\frac{P}{N\left(\frac{PT}{y}-h\right)}\right) \quad \geq \alpha \tag{S32}$$

$$E(\hat{h}) = \quad E\left(\frac{PT}{y} - \frac{P}{\alpha N}\right) \quad \geq h\,. \tag{S33}$$

***Arditi-Akçakaya***

$$\lambda = F(N,P,T,\theta) = \frac{\alpha N}{P^m + \alpha h N}PT \tag{S34}$$

$$\ln \mathcal{L}(\theta|y) = -n\frac{\alpha NPT}{P^m + \alpha h N} + \ln \frac{\alpha NPT}{P^m + \alpha h N}\sum_{i=1}^{n} y_i - \ln\left(\prod_{i=1}^{n} y_i!\right) \tag{S35}$$

$$\frac{d\, \ln \mathcal{L}(\theta|y)}{d\alpha} = 0 \implies \hat{\alpha} = \frac{P^m \sum_{i=1}^{n} y_i}{nNPT - hN\sum_{i=1}^{n} y_i} = \frac{\bar{y}P^m}{N\,(PT - h\bar{y})} \tag{S36}$$

$$\frac{d\, \ln \mathcal{L}(\theta|y)}{dh} = 0 \implies \hat{h} = \frac{nPT}{\sum_{i=1}^{n} y_i} - \frac{P^m}{\alpha N} = \frac{PT}{\bar{y}} - \frac{P^m}{\alpha N} \tag{S37}$$

$$\frac{d\, \ln \mathcal{L}(\theta|y)}{dm} = 0 \implies \hat{m} = \frac{\ln\left(\alpha N\left(\frac{PT}{\bar{y}} - h\right)\right)}{\ln P} \tag{S38}$$

Note that $\frac{\alpha NPT}{\bar{y}} \geq \alpha N h$ because $\frac{PT}{\bar{y}} \geq h$ (that is, $\frac{\bar{y}}{PT} \leq \frac{1}{h}$) because the mean number of prey eaten per predator per time of the experiment cannot exceed the maximum possible number of prey eaten per predator in that time, $1/h$. Therefore $\hat{m}$ is always a positive number.

As above, we can use the facts that $\hat{a}$ and $\hat{h}$ are *convex* functions of $y$ to show that there is positive bias in both. We check for bias in $\hat{m}$ with

$$E(\hat{m}) \quad = \quad \sum_{y=0}^{\infty} \hat{m}\frac{\lambda^y e^{-\lambda}}{y!} \tag{S39}$$

$$= \quad \sum_{y=0}^{\infty}\left(\frac{\ln(\alpha N) + \ln\left(\frac{PT}{y} - h\right)}{\ln P}\right)\frac{\lambda^y e^{-\lambda}}{y!}\,. \tag{S40}$$

The expression for $\hat{m}$ is a *convex* function of $y$. Jensen's inequality therefore dictates that

$$E\left(\frac{\ln(\alpha N) + \ln\left(\frac{PT}{y} - h\right)}{\ln P}\right) \geq \frac{\ln(\alpha N) + \ln\left(\frac{PT}{E(y)} - h\right)}{\ln P}\,. \tag{S41}$$

Substituting $E(y) = \lambda = F(N, P, T, \theta)$ into this expression we find that

$$E(\hat{m}) \geq \frac{\ln(\alpha N) + \ln\left(\frac{PT}{\lambda} - h\right)}{\ln P} = m\,, \tag{S42}$$

which implies that the bias on $\hat{m}$ will always be positive.

#### S7.2 Binomial

The binomial distribution has a single parameter, $p$, specifying the probability of a "successful" event. The expected number of successful events in $N$ Bernoulli trials is $Np$ with variance $Np(1-p)$. We express the likelihood function as being for $n$ replicates of such a binomial process (i.e. $nN$ Bernoulli trials). The expected total number of successes is therefore $nNp$ with variance $nNp(1-p)$, and $y_i$ refers to the number of successes in the $i$th replicate.

$$f(y|p) = \binom{N}{y} p^y (1-p)^{N-y} \qquad \text{Probability function} \tag{S43}$$

$$\mathcal{L}(p|y) = \prod_{i=1}^{n} f(y_i) = \prod_{i=1}^{n} \frac{N!}{y_i!(N-y_i)!} p^{y_i}(1-p)^{N-y_i} \qquad \text{Likelihood function} \tag{S44}$$

$$\ln \mathcal{L}(p|y) = \sum_{i=1}^{n} \ln\left(\frac{N!}{y_i!(N-y_i)!}\right) + \sum_{i=1}^{n} y_i \ln(p) + \left(nN - \sum_{i=1}^{n} y_i\right) \ln(1-p) \qquad \text{log-Likelihood} \tag{S45}$$

$$\frac{d\ \ln \mathcal{L}(p|y)}{dp} = (1-p)\sum_{i=1}^{n} y_i - p\left(Nn - \sum_{i=1}^{n} y_i\right) = 0 \tag{S46}$$

$$\implies \hat{p} = \frac{\sum_{i=1}^{n} y_i}{nN} = \frac{\bar{y}}{N} \qquad \text{MLE} \tag{S47}$$

The Binomial is used in the context of functional-response experiments where eaten prey are not replaced. Given an initial level of $N$ prey available and a presumed functional response, $F(N, P, \theta)$, the predicted $k$ number of prey consumed (i.e. "successful" events) is determined by integrating the dynamical equation

$$\int_{N}^{N-k} \frac{dN}{-F(N, P, \theta)} = \int_{0}^{T} dt \tag{S48}$$

and solving for $k$ (see also Rosenbaum & Rall, 2018). Note that we here abuse our own notation for the functional response $F$ because the time period of the experiment is explicitly considered in the right hand side of the equation (i.e. $\int_0^T dt = T$). For many functional responses, the analytical solution of this integral leads to a transcendental equation for $k$ of the form $ke^k = x$. In these cases, the solution(s) for $k$ can be determined with the Lambert $W$ function (Lehtonen, 2016), which we will write below as $W_0(x)$. Upon determining $k$, we can compute the likelihood given the functional-response parameters using $p = \frac{k}{N}$, with $y_i$ referring to the number of eaten prey in the $i$th replicate.

***Holling Type I***
For the Holling Type I, the integration of eqn. S48 does not lead to a transcendental equation, hence the solution does not necessitate use of the Lambert $W$ function.

$$p = \frac{k}{N} = 1 - e^{-aPT} \tag{S49}$$

$$\ln \mathcal{L}(\theta|y) = \sum_{i=1}^{n} \ln \binom{N}{y_i} + \sum_{i=1}^{n} y_i \ln(1 - e^{-aPT}) + (N - \sum_{i=1}^{n} y_i) \ln(e^{-aPT}) \tag{S50}$$

$$\frac{d \ \ln \mathcal{L}(\theta|y)}{da} = 0 \implies \hat{a} = \frac{\ln\left(\frac{N}{N-\bar{y}}\right)}{PT} \tag{S51}$$

We check for bias on $\hat{a}$ with

$$E(\hat{a}) \ = \ \sum_{y=0}^{\infty} \hat{a} \binom{N}{y} p^y (1-p)^{N-y} \tag{S52}$$

$$= \ \sum_{y=0}^{\infty} \left( \frac{\ln\left(\frac{N}{N-y}\right)}{PT} \right) \binom{N}{y} p^y (1-p)^{N-y} . \tag{S53}$$

The expression for $\hat{a}$ is a *convex* function of $y$. Jensen's inequality therefore dictates that

$$E\left( \frac{\ln\left(\frac{N}{N-y}\right)}{PT} \right) \geq \frac{\ln\left(\frac{N}{N-E(y)}\right)}{PT} . \tag{S54}$$

Substituting $E(y) = k$ into this expression we find that

$$E(\hat{a}) \geq \frac{\ln\left(\frac{N}{N-(N-Ne^{-aPT})}\right)}{PT} = a , \tag{S55}$$

meaning that there is positive bias on $\hat{a}$ (unlike for experiments in which eaten prey are continually replaced).

***Holling Type II***

$$p = \frac{k}{N} = 1 - \frac{1}{ahN} W_0 \left[ ahNe^{-a(PT-hN)} \right] \tag{S56}$$

$$\frac{d \ \ln \mathcal{L}(\theta|y)}{da} = 0 \implies \hat{a} = \frac{\ln\left(\frac{N}{N-\bar{y}}\right)}{PT - h\bar{y}} \tag{S57}$$

$$\frac{d \ \ln \mathcal{L}(\theta|y)}{dh} = 0 \implies \hat{h} = \frac{aPT + \ln\left(\frac{N-\bar{y}}{N}\right)}{a\bar{y}} = \frac{PT}{\bar{y}} + \frac{\ln\left(\frac{N-\bar{y}}{N}\right)}{a\bar{y}} \tag{S58}$$

As above, we can use the fact that $\hat{a}$ is a *convex* function of $y$ to show that there is positive bias on $\hat{a}$. The bias of $\hat{h}$ is a more nuanced, depending on the dominance of its first and second terms. $\hat{h}$ is a *convex* function of $y$, and hence biased positive, when $\bar{y} << N$, much like under a Poisson (i.e. replacement) scenario. $\hat{h}$ is a *concave* function of $y$ when $\frac{d\hat{h}}{dy} > 0$, but this only occurs as $\bar{y}$ approaches $N$ within the domain where $\hat{h}$ remains positive. Hence, negative bias will only occur when significant depletion occurs.

***Arditi-Akçakaya***

$$p = \frac{k}{N} = 1 - \frac{P^m}{ahN} W_0 \left[ \frac{ah}{P^m} N e^{-\frac{a}{P^m}(PT - hN)} \right] \tag{S59}$$

$$\frac{d \ \ln \mathcal{L}(\theta|y)}{da} = 0 \implies \hat{a} = \frac{P^m \ln\left(\frac{N}{N-\bar{y}}\right)}{PT - h\bar{y}} \tag{S60}$$

$$\frac{d \ \ln \mathcal{L}(\theta|y)}{dh} = 0 \implies \hat{h} = \frac{aPT + P^m \ln\left(\frac{N-\bar{y}}{N}\right)}{a\bar{y}} = \frac{PT}{\bar{y}} + \frac{P^m \ln\left(\frac{N-\bar{y}}{N}\right)}{a\bar{y}} \tag{S61}$$

$$\frac{d \ \ln \mathcal{L}(\theta|y)}{dm} = 0 \implies \hat{m} = \frac{\ln\left(\frac{a(PT-h\bar{y})}{\ln(\frac{N}{N-\bar{y}})}\right)}{\ln P} \tag{S62}$$

As above, we can use the fact that $\hat{a}$ is a *convex* function of $y$ to show that there is positive bias on $\hat{a}$. Similarly, there is positive bias on $\hat{m}$ since it is a *convex* function of $y$. As for the Holling Type II response, the bias of $\hat{h}$ is a more nuanced, but will be positive unless significant depletion occurs (i.e. $\bar{y}$ approaches $N$).

##### S7.3 Normal and log-Normal

The Normal (Gaussian) distribution has two parameters, with $\mu$ specifying the expected number of events and $\sigma^2$ their variance.

$$f(y|\mu,\sigma) = \frac{1}{\sqrt{2\pi\sigma^2}} e^{\frac{-(y-\mu)^2}{2\sigma^2}} \quad \text{Probability function} \tag{S63}$$

$$\mathcal{L}(\mu,\sigma|y) = \prod_{i=1}^{n} f(y_i) = \prod_{i=1}^{n} \frac{1}{\sqrt{2\pi\sigma^2}} e^{\frac{-(y_i-\mu)^2}{2\sigma^2}} \quad \text{Likelihood function} \tag{S64}$$

$$= \frac{1}{\left(\sqrt{2\pi\sigma^2}\right)^n} e^{\frac{-\sum_{i=1}^{n}(y_i-\mu)^2}{2\sigma^2}} \tag{S65}$$

$$\ln \mathcal{L}(\mu,\sigma|y) = -\frac{1}{2} n \ln\left(2\pi\sigma^2\right) - \frac{1}{2\sigma^2}\sum_{i=1}^{n}(y_i-\mu)^2 \quad \text{log-Likelihood} \tag{S66}$$

$$= -\frac{1}{2} n \ln\left(2\pi\sigma^2\right) - \frac{\sum_{i=1}^{n}(y_i)^2}{2\sigma^2} - \frac{n\mu^2}{2\sigma^2} + \frac{2\mu\sum_{i=1}^{n} y_i}{2\sigma^2} \tag{S67}$$

$$\frac{d\ \ln \mathcal{L}(\mu,\sigma|y)}{d\mu} = \frac{1}{\sigma^2}\sum_{i=1}^{n}(y_i-\mu) = \sum_{i=1}^{n} y_i - n\mu = 0 \tag{S68}$$

$$\implies \hat{\mu} = \frac{1}{n}\sum_{i=1}^{n} y_i = \bar{y} \quad \text{MLE} \tag{S69}$$

Given a presumed functional response, $F(N,P,T,\theta)$, and an experiment in which eaten prey are continually replaced, we therefore have

$$\ln \mathcal{L}(\theta|y) = -\frac{1}{2} n \ln\left(2\pi\sigma^2\right) - \frac{1}{2\sigma^2}\sum_{i=1}^{n}(y_i - F(N,P,T,\theta))^2. \tag{S70}$$

Using the same derivation steps as for the Poisson, we can show that all MLEs assuming a Normal distribution are the same as those derived under the Poisson. For brevity, we show this for only the Arditi-Akçakaya functional response.

***Arditi-Akçakaya***

$$F(N,P,T,\theta) = \frac{\alpha N}{P^m + \alpha h N} PT \tag{S71}$$

$$\ln \mathcal{L}(\theta) = -\frac{1}{2} n \log\left(2\pi\sigma^2\right) - \frac{1}{2\sigma^2} \sum_{i=1}^{n} \left( y_i - \frac{\alpha NPT}{P^m + \alpha h N} \right)^2 \tag{S72}$$

$$\frac{d\ \ln \mathcal{L}(\theta)}{d\alpha} = \frac{NTP^{m+1}\left(n\alpha NPT - \sum_{i=1}^{n} y_i \left(P^m + \alpha h N\right)\right)}{\sigma^2 \left(P^m + \alpha h N\right)^3} = 0 \tag{S73}$$

$$\implies \hat{\alpha} = \frac{P^m}{N\left(\frac{PT}{\bar{y}} - h\right)} \tag{S74}$$

$$\frac{d\ \ln \mathcal{L}(\theta)}{dh} = \frac{\alpha^2 N^2 PT\left(n\alpha NPT - \sum_{i=1}^{n} y_i \left(P^m + \alpha h N\right)\right)}{\sigma^2 \left(P^m + \alpha h N\right)^3} = 0 \tag{S75}$$

$$\implies \hat{h} = \frac{PT}{\bar{y}} - \frac{P^m}{\alpha N} \tag{S76}$$

$$\frac{d\ \ln \mathcal{L}(\theta)}{dm} = \frac{\alpha NTP^{m+1} \ln P\left(n\alpha NPT - \sum_{i=1}^{n} y_i \left(P^m + \alpha h N\right)\right)}{\sigma^2 \left(P^m + \alpha h N\right)^3} = 0 \tag{S77}$$

$$\implies \hat{m} = \frac{\ln\left(\alpha N\left(\frac{PT}{\bar{y}} - h\right)\right)}{\ln P} \tag{S78}$$

These MLEs are the same as those derived when assuming a Poisson process and are thereby subject to the same qualitative biases. (As discussed in Section S6, they may also suffer from addition bias associated with the estimation of the "nuisance" parameter $\sigma$.)

Under the assumption that the numbers of prey eaten are random variables drawn from a log-Normal distribution, the likelihood is written as

$$\mathcal{L}(\mu, \sigma | y) = \prod_{i=1}^{n} \frac{1}{y_i\sqrt{2\pi\sigma^2}} e^{\frac{-(\ln y_i - \mu)^2}{2\sigma^2}}. \tag{S79}$$

Functional response parameter MLEs derived assuming a log-Normal distribution are therefore equivalent to those as derived under the Normal likelihood except that the observations $y_i$ are each log-transformed. This causes the MLEs derived under the log-Normal likelihood to become functions of the geometric mean number of prey eaten, rather than the arithmetic mean, thereby introducing additional bias due to an added layer of nonlinearity.
